# Deconvolving the spatiotemporal chromatin landscape through the cell cycle

**DOI:** 10.64898/2026.07.07.731387

**Authors:** Trung Q. Tran, Yulong Li, David M. MacAlpine, Alexander J. Hartemink

**Affiliations:** Department of Computer Science, Duke University, Durham, NC 27708, USA; Department of Pharmacology and Cancer Biology, Duke University Medical Center, Durham, NC 27710, USA

## Abstract

Profiling genomic processes during the cell cycle is challenging because synchronized populations gradually lose synchrony as individual cells progress through the cycle at different rates and divide asymmetrically. Researchers have addressed this challenge by modeling the loss of synchrony and applying sophisticated branching process deconvolution methods to mitigate the effects of imperfect synchrony. Such methods have been used to deconvolve cell cycle transcription, but despite the central role of the chromatin landscape in orchestrating eukaryotic transcription and replication, comparable approaches to deconvolving cell cycle chromatin occupancy have not been developed, because the data are orders of magnitude larger and because DNA replication introduces non-uniform copy number effects across the genome during S phase. We present CyCLOPS, a computational framework that overcomes these technical challenges, enabling deconvolution of the genome-wide chromatin landscape throughout the cell cycle at high spatiotemporal resolution. We apply CyCLOPS to MNase-seq data collected from synchronized yeast populations at 10-minute intervals to produce the first dynamic atlas of genome-wide chromatin occupancy through the cell cycle, profiled at sub-minute resolution. We identify functional groups of cell cycle genes through chromatin-based clustering and uncover chromatin regulatory dynamics, including at non-genic loci. Our atlas reveals that chromatin occupancy and transcription fluctuate largely independently.

## Introduction

The cell division cycle is a fundamental process, highly conserved across all eukaryotic organisms, that governs how cells grow, replicate, and divide. Progression through the cell cycle requires precise coordination of multiple interacting cellular processes, including many that take place at the genomic level where the chromatin landscape plays a key role in the regulation of transcription and replication. Revealing how the chromatin landscape is linked to transcription and replication through the cell cycle is crucial for understanding both normal cellular function and pathological states like cancer.

In the study of the eukaryotic cell cycle, the budding yeast *S. cerevisiae* is an established model organism due to its genetic tractability and conserved cell cycle mechanisms (Hartwell and Unger 1977). Monitoring individual yeast cells as they progress through the cell cycle can be accomplished with microscopy or emerging single-cell assays. Live-cell microscopy enables longitudinal tracking of the progression of individual cells and provides spatial information at the cellular scale, but remains technically challenging, requiring multidisciplinary expertise with limited standardized approaches (Cuny *et al*. 2022). Moreover, microscopy can only monitor a few predetermined molecules at a time, so monitoring larger numbers of molecules requires large-scale parallel experiments (Litsios *et al*. 2019). In contrast, single-cell sequencing-based assays enable the simultaneous quantification of thousands of molecules, but need to estimate the cell cycle time of each cell indirectly (Guo and Chen 2024), and suffer from data sparsity issues that often necessitate aggregation back into bulk analyses (Gasch *et al*. 2017; Nadal-Ribelles *et al*. 2019). Given these constraints, the most common approach for high-throughput molecular quantification during the yeast cell cycle is to profile populations of cells that have been synchronized to progress simultaneously through the cycle in a coordinated fashion (Juanes 2017).

High-throughput assays of synchronized populations of cells have proved exceedingly fruitful over the years. Studies have systematically profiled cell cycle transcript levels (Cho *et al*. 1998; Granovskaia *et al*. 2010; Pramila *et al*. 2006; Spellman *et al*. 1998) and protein abundance (Flory *et al*. 2006), as well as the two together (Kelliher *et al*. 2018) and in conjunction with metabolites (Campbell *et al*. 2020). In recent years, studies have also monitored chromatin state during the cell cycle; for instance, the dynamic nucleosome architecture at transcription start sites (Deniz *et al*. 2016) and chromatin occupancy at origins of replication (Li *et al*. 2021) and at gene bodies (Li *et al*. 2026) have been assayed throughout the cell cycle, while Santos *et al*. (2022) explored how the process of replication influences nucleosome organization and transcription factor binding.

However, accurately interpreting measurements from synchronized populations of cells requires addressing fundamental technical challenges that are inherent to synchronization. First, perfect synchrony is neither attainable nor maintainable: Cells exhibit inherent variability at the time of release and further lose synchrony as they progress through the cell cycle at different rates. Second, the use of chemicals to induce synchrony can alter cellular state: Cells accumulate protein and mass during arrest, resulting in larger-than-normal cells with perturbed physiology upon release (Rosebrock 2017). Third, asymmetric cell division adds a further layer of complexity (Sunchu and Cabernard 2020): In budding yeast, for example, daughter cells emerge smaller than mother cells, requiring extended time for growth in G1 (Hartwell and Unger 1977; Lord and Wheals 1980).

Researchers have addressed some of these challenges by applying temporal deconvolution methods to transcriptional data from populations of yeast cells (Bar-Joseph *et al*. 2004; Guo *et al*. 2013; Orlando *et al*. 2007, 2009; Qiu *et al*. 2006; Rowicka *et al*. 2007; Siegal-Gaskins *et al*. 2009), but no comparable method has been developed for chromatin occupancy data because it poses two unique analytical challenges. First, chromatin occupancy assays generate massive two-dimensional occupancy profiles across the entire genome, necessitating orders of magnitude more computation. Second, during S phase, different regions of the genome replicate at different times, creating variable DNA copy number effects across the genome that complicate the interpretation of chromatin occupancy data, in contrast to transcript, protein, or metabolite data.

Here we present CyCLOPS (Cycling Chromatin Landscape Occupancy Profiling System), a temporal deconvolution framework to profile the dynamics of the genome-wide chromatin landscape throughout the cell cycle. CyCLOPS deconvolves data from synchronized populations of cells to generate profiles of chromatin occupancy and transcription at high temporal resolution, while simultaneously generating replication timing profiles to correct for non-uniform copy number effects. By applying CyCLOPS to yeast MNase-seq data, we provide the first dynamic atlas of a eukaryotic chromatin occupancy landscape through the cell cycle genome-wide, including at genes and replication origins. We identify specific functional gene groups with coordinated cyclic chromatin-transcriptional dynamics and find patterns of cyclic +1 nucleosomes linked to cell cycle gene expression. Beyond genic regions, our framework reveals cell cycle dynamics in antisense and non-coding regions, and detects mother/daughter-specific chromatin differences. Surprisingly, we find that chromatin occupancy and gene expression cycle largely independently. CyCLOPS is a powerful approach for revealing the cell cycle chromatin landscape at unprecedented spatiotemporal resolution.

## Results

### CyCLOPS enables genome-wide deconvolution of chromatin dynamics throughout the cell cycle

We developed CyCLOPS (Cycling Chromatin Landscape Occupancy Profiling System), a framework to deconvolve both transcription and chromatin genome-wide throughout the cell cycle into average, idealized single-cell profiles that correspond to portions of a branching process model (Fig. 1a). The cell cycle branching process is characterized by various parameters that are estimated from flow cytometry data (Supplemental Figure S1) by a model called CLOCCS (Characterizing Loss of Cell Cycle Synchrony) (Orlando *et al*. 2009) (Supplemental Figure S2).

**Figure 1.**
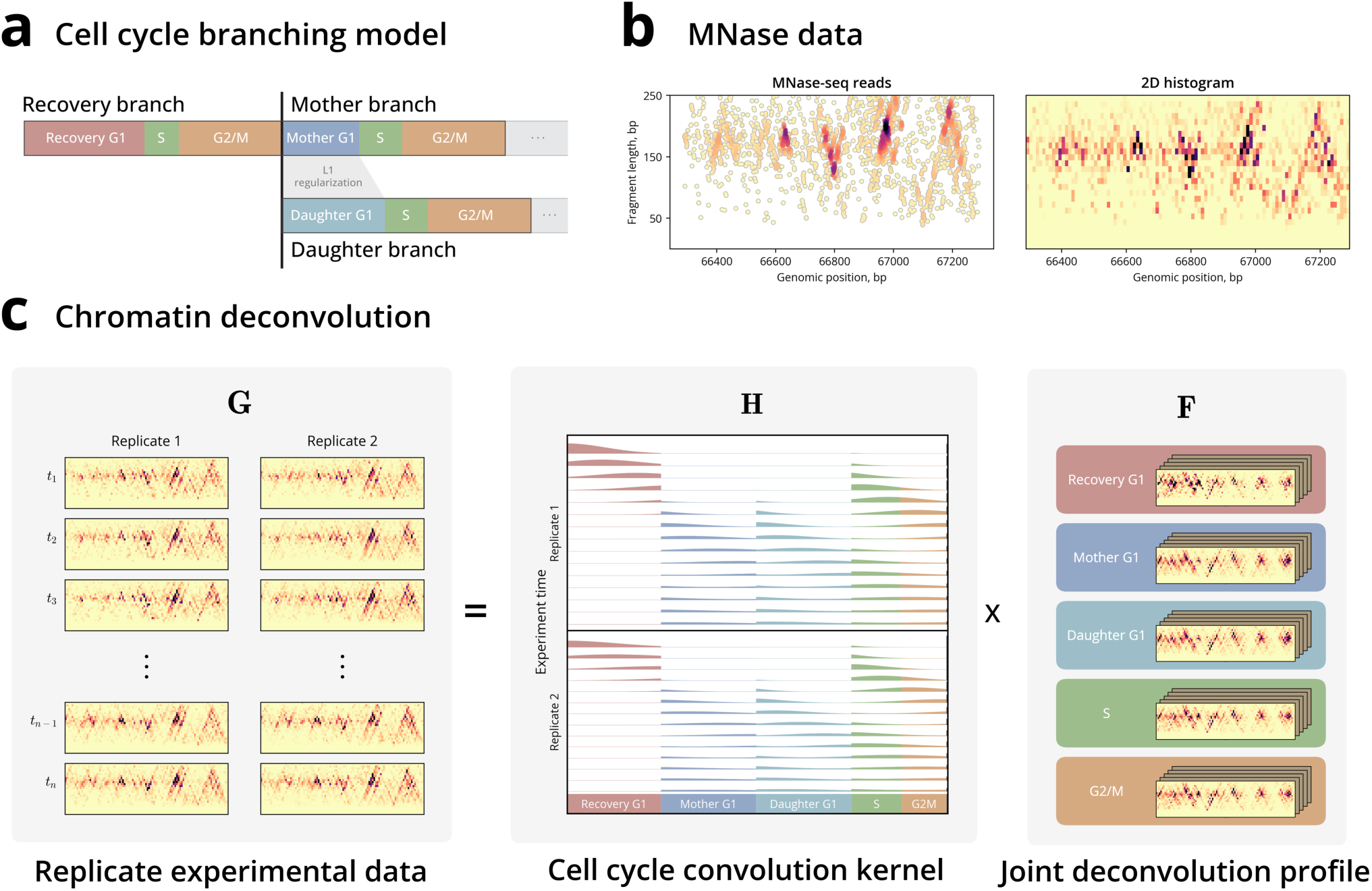
CyCLOPS deconvolves chromatin data into phase-specific single-cell profiles. **(a)** The cell cycle can be modeled as a branching process. An initial cohort of cells grows and branches into distinct mother and daughter cohorts. Mother G1 and daughter G1 intervals are regularized for similarity. **(b)** Left: Raw chromatin data are depicted as midpoint positions along the x-axis and fragment lengths on the y-axis, colored by density of nearby reads. Right: The same data are binned into a fixed-size histogram for deconvolution. **(c)** Population-level chromatin data are modeled as a convolution, or mixture, of idealized single-cell chromatin data, which we seek to estimate. Left: Raw, experimental chromatin data are binned at each time point in each replicate experiment. Middle: Cell cycle parameters are encoded in stacked convolution kernels where each experimental time point in each replicate (row) contains a mixture of cells at each deconvolved time point (column) of the idealized cell cycle. Right: Deconvolved profiles represent chromatin occupancy at high temporal resolution that can be assigned precisely to each phase of the cell cycle.

We incorporate these CLOCCS-derived parameter estimates into CyCLOPS to deconvolve data reflecting the chromatin state. To generate our chromatin data set, we transform the raw variable fragment-length chromatin occupancy values into genome-wide two-dimensional histograms (Fig. 1b), segmenting the entire 12-million base pair yeast genome across 250 fragment lengths into 30 million spatial bins.

Beyond correcting for population heterogeneity, chromatin occupancy data requires an additional correction specific to genomic binding assays: copy number correction. During S phase, DNA replication creates a varying copy number effect as portions of the genome replicate at different times. We describe our approach to correcting this effect in the next section.

Finally, CyCLOPS deconvolves the genomic assays by jointly modeling independent experimental time courses to generate unified deconvolution profiles for both the chromatin (Fig. 1c) and for gene expression. Our model enhances temporal resolution from 15 time points collected at 10-minute intervals to 257 time points at sub-minute intervals. CyCLOPS is the first genome-wide deconvolution of the chromatin landscape, incorporating both fine-resolution spatial binning and copy number correction.

### Deconvolution of MNase-seq generates high-quality replication profiles enabling correction of pervasive copy number variation

Unlike studies of cell cycle gene expression, chromatin analysis faces the additional challenge of distinguishing true chromatin changes from copy number effects created by the gradual replication of DNA over the course of S phase (Fig. 2a). Within CyCLOPS, we developed a technique to correct copy number effects for more accurate analysis of true chromatin signals during S phase (Fig. 2b).

**Figure 2.**
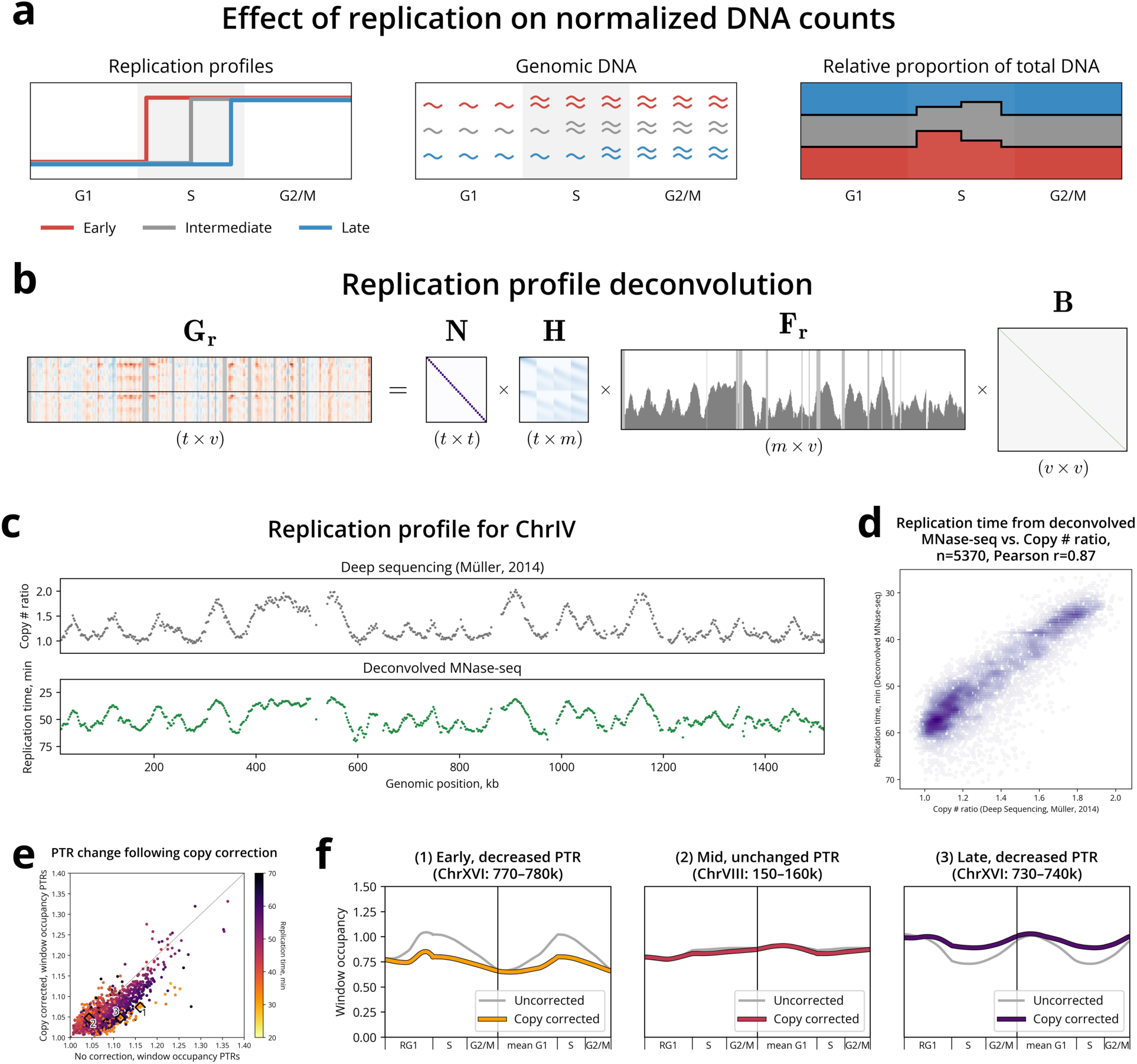
MNase-seq-derived replication profiles enable copy number correction. **(a)** Cartoon illustrating how replication creates a differential proportion of read counts at regions replicating early vs. late. Left: Replication profile curves for early (red), mid (gray), and late (blue) replication regions. Middle: Read count depiction of total DNA for these regions. Right: When normalized to equal total read counts for each time point, counts in early regions are inflated during S-phase replication; conversely, reads in late regions are deflated during this period. **(b)** Replication profile deconvolves the genome-wide read counts **G***_r_* into diagonal normalization matrix **N**, convolution kernel **H**, replication profile **F***_r_*, and diagonal baseline occupancy matrix **B**. **(c)** Chromosome IV shows consistent timing profiles between those collected from deep sequencing copy ratios from Müller *et al*. (2014) (top) and our deconvolved MNase-seq timing profile (bottom). **(d)** These two approaches are strongly correlated (Pearson r=0.87). **(e)** Copy number correction systematically decreases PTR across most genomic regions (points below diagonal), as expected. Notable examples are highlighted as numbered diamonds. **(f)** These examples depict three distinct effects of copy correction: (1) Early replicating region shows decreased PTR, correcting for artificially inflated S-phase signals; (2) Mid-replicating region exhibits minimal PTR change; (3) Late replicating region shows decreased PTR, correcting for artificially deflated signals when early regions have already replicated.

Our approach profiles replication timing using large 10 kb averages of MNase-seq occupancy. Our replication timing profiles agreed with established deep-sequencing-derived profiles from Müller *et al*. (2014) (Fig. 2c,d), with a Pearson correlation of *r* = 0.87. Our profiles were also consistent with 437 established origins annotated as early or late replicating by Belsky *et al*. (2015), showing separation between early and late annotated origins along our estimated replication timing (Supplemental Figure S4a).

We refined this origin set to identify high-confidence firing origins suitable for downstream analysis. We filtered the 437 origins using two criteria: (1) inferred firing, where origins showed earlier replication timing than their neighboring regions, and (2) at least 5 kb away from other origins to ensure selected origins were not replicated by nearby replication forks. These 49 “inferred-firing” origins showed statistically significant enrichment for separating early and late replication classes (p*<*0.0001), whereas the full set did not (p=0.169) (Supplemental Figure S4b). We use this filtered set of inferred-firing origins for subsequent chromatin analysis at replication origins.

Having validated our replication profiles against both established replication timing and annotated origins, we incorporated our profiles directly into CyCLOPS to correct for copy number effects. This correction reveals the deconvolved chromatin profiles linked to true regulatory dynamics rather than replication-driven artifacts.

To evaluate the effect of this correction, we computed peak-to-trough ratios (PTRs), calculated as the ratio of the 90th percentile over 10th percentile occupancy of all reads within each 10 kb window across the genome. We observe a global decrease in PTR after copy number correction, with most loci falling below the diagonal in the PTR comparison plot (Fig. 2e). This shows that, without correction, copy number variation pervasively inflates apparent chromatin cycling signals. Within this global trend, we identified two distinct effects: systematic PTR decreases linked to replication timing, and a subset of windows where correction reveals previously masked cell cycling chromatin dynamics.

Within this global pattern, we observed a clear relationship between regions with decreased PTR and replication timing. The earliest replicating regions show the largest decrease in PTR (Fig. 2f(1)), correcting for inflated read counts in early S phase. The latest replicating regions exhibit a gradient of decreases, from minimal (Fig. 2f(2)) to distinct decreases in PTR (Fig. 2f(3)). For these late replicating regions, prior to applying copy correction, read counts are artificially deflated when earlier regions have replicated. These patterns demonstrate the necessity for copy number correction prior to downstream analyses.

### CyCLOPS deconvolution improves the temporal and cell-cycle-phase resolution of chromatin signal

Using CyCLOPS, we deconvolved the transcriptome and chromatin genome-wide. Deconvolution enhances detection of true cell cycle signal while removing artifacts from copy number doubling and biological replicate variability. We demonstrate this improved signal at both the locus level and on a genomic scale.

We first compared the deconvolved (Fig. 3) and raw data (Supplemental Figure S5) at a well-characterized locus. While both capture cell cycle chromatin changes, deconvolution substantially improves cell cycle phase identification, temporal resolution (10 min into sub-minute), and signal-to-noise by jointly modeling the replicate experiments. Within this locus, we recover regulatory dynamics of known cell cycle regulators.

**Figure 3.**
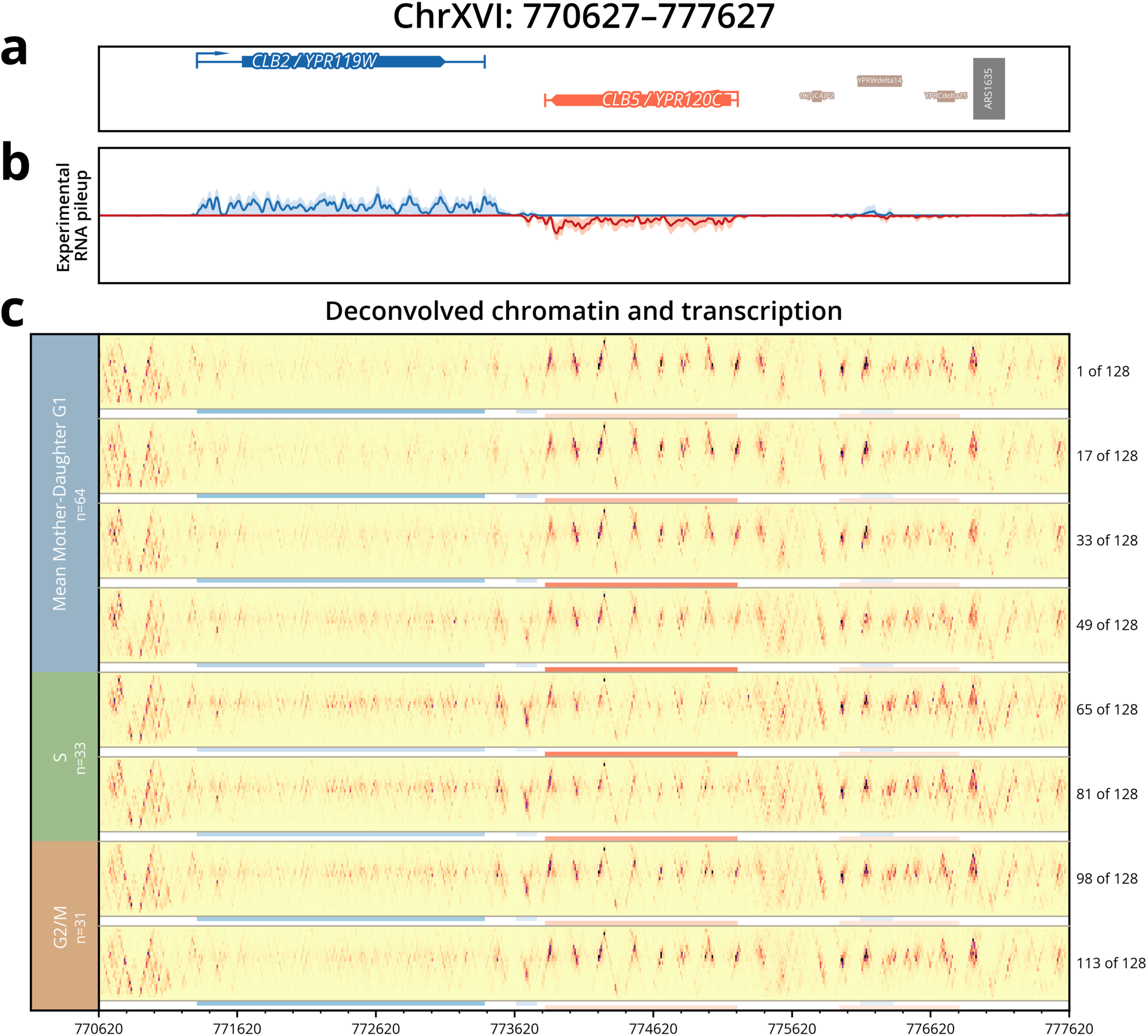
CyCLOPS enables high resolution analysis of chromatin and transcription across the genome. Locus plot shows the region surrounding cyclin genes *CLB2* and *CLB5*. **(a)** Genomic context shows annotations of genes and origin of replication. **(b)** Average stranded RNA-seq pileup is shown as thin blue (Watson) and red (Crick) trace curves. The 5th–95th percentiles across time points are depicted as light red and blue envelopes (wider envelopes indicate wider variation in expression over the time course). **(c)** The deconvolved chromatin and transcription profiles are plotted as heatmaps and colored lines. Of the 128 deconvolved time points for the presented branch, 8 are presented for brevity. Mother and daughter G1 profiles are averaged to focus analyses on concordant signal. Transcript levels for corresponding genes are depicted as shades of blue or red, with darker values indicating higher expression. The gene body nucleosomes for *CLB2* increase in occupancy from the end of G1 through S phase coinciding with its lower expression level. *CLB5*, on the other hand, exhibits greater disorganization and increased expression level in this same period. Upstream (to the right) of *CLB5*, early and efficient origin *ARS1635* shows a clear footprint signal at the end of G1 and the start of S phase.

The region contains cell cycle regulated B-type cyclin genes *CLB2* and *CLB5* and the early firing origin ARS1635. These B-type cyclins have distinct functional timing: *CLB2* controls G2/M progression and is primarily expressed in late G2/early G1, while *CLB5* drives replication initiation and is most actively transcribed during late G1/S. During gene transcription, nucleosomes shift and can even be evicted from the gene body to allow RNA polymerase (Pol II) to bind and transcribe (Kulaeva *et al*. 2010). During late G1/S, we observed chromatin patterns matching this polymerase activity: *CLB2* shows high gene body nucleosome occupancy reflecting transcriptional inactivity, whereas *CLB5* exhibits the nucleosome disorganization that is a hallmark of active transcription. Meanwhile, the nearby early firing origin *ARS1635* displays a protected region whose footprint is dynamic across the cell cycle, with peak protection at the G1/S border—consistent with pre-replicative complex (pre-RC) formation.

This signal is consistent with the expected timing of replication in S phase. As a control, the nearby stably transcribed gene *THI22* (Supplemental Figure S6) displays relatively unchanging chromatin signal across all cell cycle phases.

To summarize the broad chromatin changes revealed by our genome-wide deconvolution, we defined three measures of genic chromatin state: promoter occupancy (small fragments bound to gene promoters), nucleosome entropy (disorganization of nucleosomes within gene bodies), and nucleosome occupancy (presence of nucleosomes in gene bodies). We use the previously defined PTR to quantify cyclicity.

We compared the change in PTR for 5,774 genes before and after deconvolution (Supplemental Figure S7). Deconvolution broadly increased PTR for nucleosome metrics, indicating improved cell cycle signals. In contrast, promoter occupancy showed mixed effects following deconvolution, with many genes decreasing in PTR. We found that small fragment signals were more susceptible to noise in the raw data. The reduction in PTR reflects the removal of false cyclicity signals.

Together, CyCLOPS enhances detection of cell cycle chromatin signals both at individual loci, where it recovers known cell cycle regulation, and genome-wide, where it systematically increases cyclic signal in nucleosome metrics.

### CyCLOPS reveals differential pre-RC loading at origins and tracks replication fork progression genome-wide

Origins of replication are specific loci where DNA replication initiates during S phase. Regulation and timing of origin firing are critical to coordinate stable DNA replication in preparation for mitosis. We sought to study the chromatin dynamics at these origins. Origins vary in firing efficiency, which depends not only on the extent of pre-replicative complex (pre-RC) assembly in G1 but also on the efficiency of its conversion to the active CMG helicase in S phase and on the surrounding chromatin environment. The pre-RC is assembled through the sequential loading of ORC, Cdc6, Cdt1, and the MCM helicase (a hetero-hexamer of the proteins Mcm2–7) onto origin DNA. Pre-RC assembly licenses the origin for activation in S phase (Costa and Diffley 2022); restricting this licensing to G1, when CDK activity is low, ensures that origins fire no more than once per cell cycle.

We examined the small fragment footprint at the 50 earliest and 50 latest firing origins in each branch, selecting origins with G1 and G2 footprints per Belsky *et al*. (2015) (Supplemental Figure S8a,b), which restricts the analysis to confidently ORC-bound origins and provides the G2 ORC-alone baseline against which the G1 pre-RC footprint can be compared. Cells arrested in *α*-factor accumulate large amounts of pre-RC, priming them for replication in this extended G1 period. These amounts exceed the pre-RC levels in both G1 and S phases for the subsequent branches (Fig. 4b). Additionally, early firing origins sustain higher pre-RC amounts than late firing origins, indicating that early origins have a greater capacity for MCM loading.

**Figure 4.**
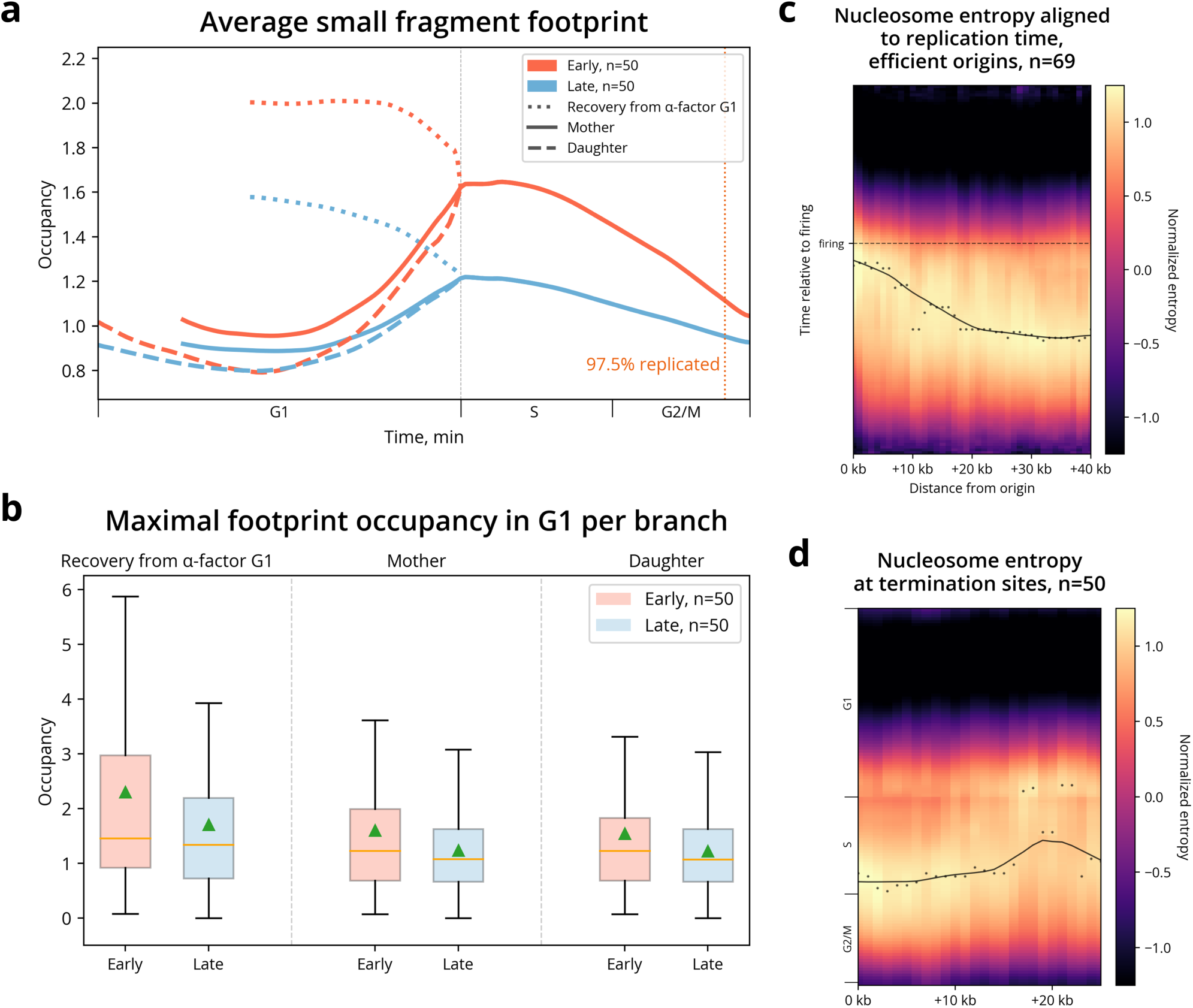
CyCLOPS reveals chromatin dynamics at origins of replication. **(a)** Average small fragment footprint timecourse for the 50 earliest and 50 latest firing origins on the recovery, mother, and daughter branches. Selected origins have G1 and G2 footprints as annotated by Belsky *et al*. (2015). Early firing origins reach higher peak subnucleosomal occupancy than late firing origins in S-phase for mother and daughter cells, consistent with greater MCM loading and higher firing efficiency. In cells recovering from *α*-factor, greater MCM accumulation in G1 reinforces the difference between early and late firing origins. The orange vertical line marks where 97.5% of the genome is replicated, based on our replication timing estimates, closer to the end of G2/M. This indicates S-phase extends substantially later than the CLOCCS estimate from DNA content and FACS suggests. **(b)** Maximal footprint occupancy per origin within G1 for recovery, mother, and daughter branches. In the recovery branch, both early and late firing origins show elevated G1 occupancy relative to mother and daughter branches, reflecting greater pre-RC accumulation during the extended G1 following *α*-factor arrest. Early origins remain more highly occupied than late origins across all three branches. Green triangles indicate the mean of each distribution. **(c)** Replication origins disrupt nucleosome organization through S phase. Nucleosome entropy is plotted as a heatmap across distances from firing origins and deconvolved cell cycle time. Entropy signal follows replication fork progression, disorganizing early in S phase closest to firing origins then progressively disrupting later in S phase within 5 kb away. The most entropic time point for each position is plotted as black scatter points with line of best fit (using LOWESS) as a black line. **(d)** Pattern of nucleosome disorganization continues to replication termination sites. The 50 termination sites are determined by the midpoint of the furthest pairs of adjacent origins. Closest to termination sites, entropy is at its highest in late S phase. Approximately 20 kb away from termination sites are the firing origins, where nucleosomes are disrupted in early S phase.

This relationship persists as cells re-enter the cell cycle. In mother cells (Fig. 4a), early firing origins reach higher peak footprint occupancy than late firing origins, with this early–late gap present throughout G1, peaking in early S phase. This difference reflects the greater efficiency of early firing origins, demonstrating that firing efficiency is linked to the extent of pre-RC loading. In daughter cells, early firing origins again reach higher peak footprint occupancy than late firing origins entering S phase. However, both early and late origins converge to an equivalent trough value in mid-G1, suggesting that, following cell division, pre-RC is transiently depleted, allowing daughter cells to progress through an extended G1 phase. Notably, although CLOCCS estimates S phase and G2/M to be of comparable length, our replication timing estimates place the completion of replication (97.5% of the genome replicated) near the end of G2/M. This indicates that S phase extends nearly to the end of the cell cycle, consistent with footprint occupancy dropping off late, once most of the genome has been replicated. Together, these results show how CyCLOPS temporally resolves footprint occupancy, linking pre-RC formation to origin firing efficiency.

Beyond characterizing pre-RC formation dynamics at origins, we sought to demonstrate how CyCLOPS amplifies replication fork progression signal. We began with the distance-based approach of Li *et al*. (2021) to analyze nucleosome entropy around the 49 previously identified, 69 (the top 20%) efficient origins and 50 termination sites, sites defined as the midpoint between the 50 pairs of origins with greatest distance between them. We used the deconvolved replication times to temporally align the firing times for each origin and amplify the entropy signal of fork-progression (Fig. 4c).

At these origins, nucleosomes were the most disorganized during early S phase at the origin itself. The timing of this disorganization shifts progressively later in S phase at greater distances from origins. At termination sites, nucleosomes disorganize late in S phase at the termination center, with early S-phase signals visible at the flanking origins approximately 20 kb away (Fig. 4d). These patterns are consistent with nucleosome disorganization tracking replication fork progression, as reported by Li *et al*. (2021). But by aligning the fork-progression signal on replication timing, CyCLOPS amplifies this signal and resolves the dynamics at a higher, sub-minute, temporal resolution Together, these results highlight CyCLOPS’s sub-minute temporal scale. We found differential pre-RC formation across early and late firing origins, and that nucleosome disorganization propagates with replication fork progression from origins through termination sites.

### Genes clustered by chromatin changes reveal patterns of cell cycle transcriptional regulation

We next sought to determine whether chromatin changes are linked to transcriptional regulation during the cell cycle. We compared PTR values for promoter occupancy, nucleosome entropy, and nucleosome occupancy to gene expression cyclicity, also measured using PTR. Each chromatin measure showed weak correlation with transcriptional cyclicity (*R*^2^ = 0.04, 0.09, and 0.08, respectively), indicating that chromatin dynamics and gene expression cycled largely independently at the global level (Fig. 5a). This finding is consistent with prior observations by Mahendrawada *et al*. (2025) and Moyung *et al*. (2026).

**Figure 5.**
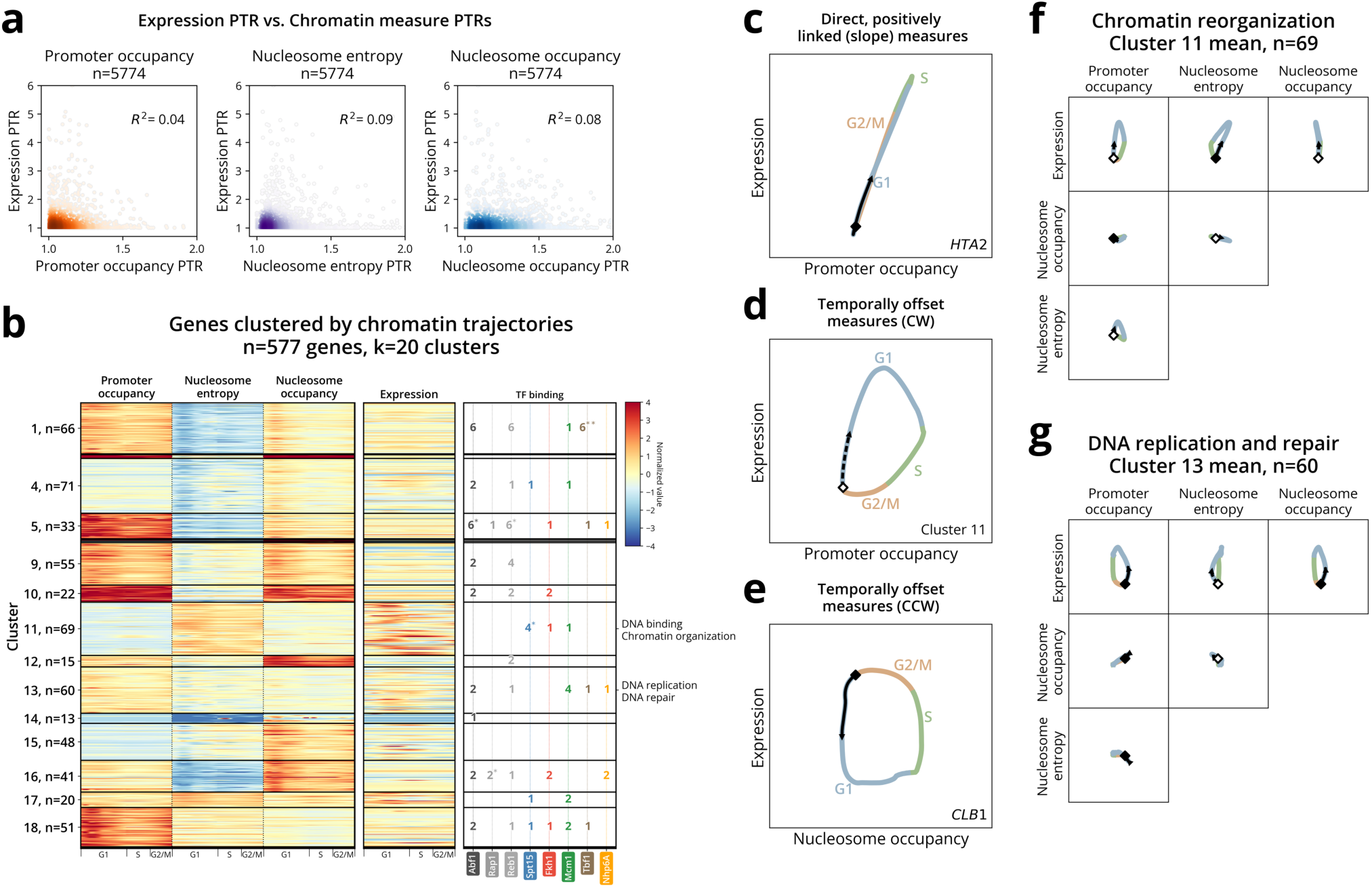
Genes clustered by chromatin changes reveal cell cycle transcription regulation patterns. **(a)** Chromatin and transcription cycle largely independently at the global level. Scatter plots compare PTR values for each chromatin measure against expression PTR across all genes (n=5774). Cyclicity of promoter occupancy, nucleosome entropy, and nucleosome occupancy each show weak correlation with expression cyclicity (*R*^2^ = 0.04, 0.09, and 0.08 respectively). **(b)** Genes clustered by chromatin trajectories reveal distinct functional groups with enriched transcription factor binding. Left: Heatmaps show z-score-normalized deconvolved time course values for promoter occupancy, nucleosome entropy, and nucleosome occupancy across 577 cyclic genes organized into *k* = 20 clusters. Middle: Associated expression values for each gene. Right: Transcription factor binding site counts with significant enrichment, computed using Fisher’s exact test with Benjamini-Hochberg correction (p-value: \**<*0.05, \*\**<*0.001). Notable clusters with functional enrichment include cluster 11 (DNA binding, chromatin organization) and cluster 13 (DNA replication and repair). **(c–e)** Schematic trajectory plots illustrate how to interpret chromatin–expression relationships through the cell cycle. Time courses are colored by cell cycle phase (G1: blue; S: green; G2/M: gold) with directional markers indicating counterclockwise (black diamond, solid line) or clockwise (white diamond, dotted line) temporal progression. **(c)** Direct, positively linked measures show linear trajectories. **(d)** Temporally offset measures with clockwise (CW) progression. **(e)** Temporally offset measures with counterclockwise (CCW) progression. **(f)** Cluster 11 mean trajectories show subtle chromatin dynamics coupled with strong cyclic expression, characteristic of DNA binding and chromatin organization genes that peak in G1. **(g)** Cluster 13, representing DNA replication and repair genes, mean trajectories also display more subtle chromatin and expression changes.

Beyond these global trends, we asked whether chromatin-transcriptional linkages existed within specific functional gene groups. We performed clustering that integrates all chromatin measures and transcription simultaneously, then analyzed the resulting clusters for enrichment of cell cycle functions and cell cycle transcription factor (TF) binding. To identify genes with coordinated chromatin-transcription dynamics, we selected 577 genes, the top 10% of genes, showing substantial cell cycle variation across the 4D measurement space (three chromatin metrics and expression). This variation was quantified using trajectory diameter, a unified measure of cyclicity across the measurement space.

Clustering these genes based solely on their chromatin trajectories—deliberately omitting expression— allowed us to test whether chromatin dynamics alone could be linked to transcriptional patterns. We identified *k* = 20 clusters using *k*-means clustering (Fig. 5b), selecting *k* using the within-cluster sum of squares (WCSS) elbow method, a heuristic for selecting the optimal number of clusters. Notably, Cluster 11 exhibited a striking cell cycle expression profile, with nearly all genes peaking sharply in G1, presenting a cluster of genes in which transcriptional timing is tightly linked to chromatin changes.

We next asked whether these clusters correspond to functionally related cell cycle processes. Gene ontology enrichment analysis revealed that two clusters showed significant functional enrichment, both with peak expression in G1. Cluster 11 was enriched for genes associated with DNA binding and chromatin organization, while cluster 13 showed enrichment for genes involved in DNA replication and repair processes. These patterns indicate that the clearest chromatin signatures of cell cycle regulated transcription appear in G1 genes. This pattern may be due to either inherently stronger chromatin-transcription coupling in G1 genes or because replication-related dynamics obscure S and G2/M chromatin-expression patterns.

To gain a better sense of how chromatin and expression coordinate in these clusters, we plotted chromatin measures against each other and against expression trajectories through the cell cycle. Trajectories may exhibit two high-level temporal modes: (1) direct, linear relationship, where measures are temporally linked (Fig. 5c) or (2) temporally offset hysteresis, looping measures (Fig. 5d). In both clusters 11 and 13, we see looping, temporally offset trajectories between promoter occupancy and expression (Fig. 5e,f). The trajectory’s directionality (clockwise or counter-clockwise) indicates the measure’s relationship with expression. For cluster 11, an increase in expression follows a decrease in promoter occupancy. This is reversed in cluster 13, where activated expression follows promoter binding. These differences reveal that cell cycle genes exhibit diverse patterns of promoter-expression coordination.

We then examined whether TF binding could explain the chromatin-transcription coordination within these clusters. We examined the prevalence of TF binding sites from Rossi *et al*. (2021) in gene promoters for each cluster and found differing enrichment patterns, linking chromatin and transcription in distinct regulatory modes.

First, clusters 1 and 5 showed enrichment for general transcription factors (Tbf1, Spt15) and pioneer factors (Abf1, Reb1), with cyclic promoter occupancy but stable transcription throughout the cell cycle. These enrichment patterns suggest a TF regulatory function that stabilizes transcriptional levels, consistent with the global pattern that cyclic chromatin is typically not associated with cyclic transcription.

Second, the chromatin organization cluster (cluster 11) showed enrichment for Spt15 binding, with strong transcriptional cycling despite subtle promoter occupancy dynamics. As a TATA-box binding protein, Spt15 primes promoters by establishing a transcriptionally poised state prior to activation (Lee *et al*. 2010). Our observations suggest that Spt15 may maintain this poised state throughout the cell cycle with relatively stable promoter occupancy, enabling transcriptional activation with minimal promoter remodeling.

Together, these results demonstrate that clustering genes by chromatin dynamics can identify functionally coherent cell cycle groups, including those revealing modes of TF regulatory function. In a subsequent section, we comprehensively explore the binding dynamics of a broad set of cell cycle transcription factors.

### Deconvolution reveals both canonical and nuanced cell cycle regulation

To explore CyCLOPS’s ability to link chromatin and expression changes, we examined well-characterized cell cycle gene families. Histone synthesis is tightly coupled with DNA replication during S phase. During replication, nucleosomes are disrupted to allow DNA polymerase access, then rapidly reassembled using both recycled and newly synthesized histones (Alabert and Groth 2012). Histone gene expression is precisely coordinated through the cell cycle, with transcription peaking during S phase when histone demand is highest (Marzluff and Duronio 2002). In our results, we see this tight coordination as consistent regulatory patterns in every histone gene (Fig. 6a). Each of these genes shows the same pattern of direct coupling between promoter occupancy changes and expression changes, peaking in S phase (Fig. 6a: linear shape of expression vs. promoter occupancy; b: small fragment enrichment and transcriptional activation in S phase).

**Figure 6.**
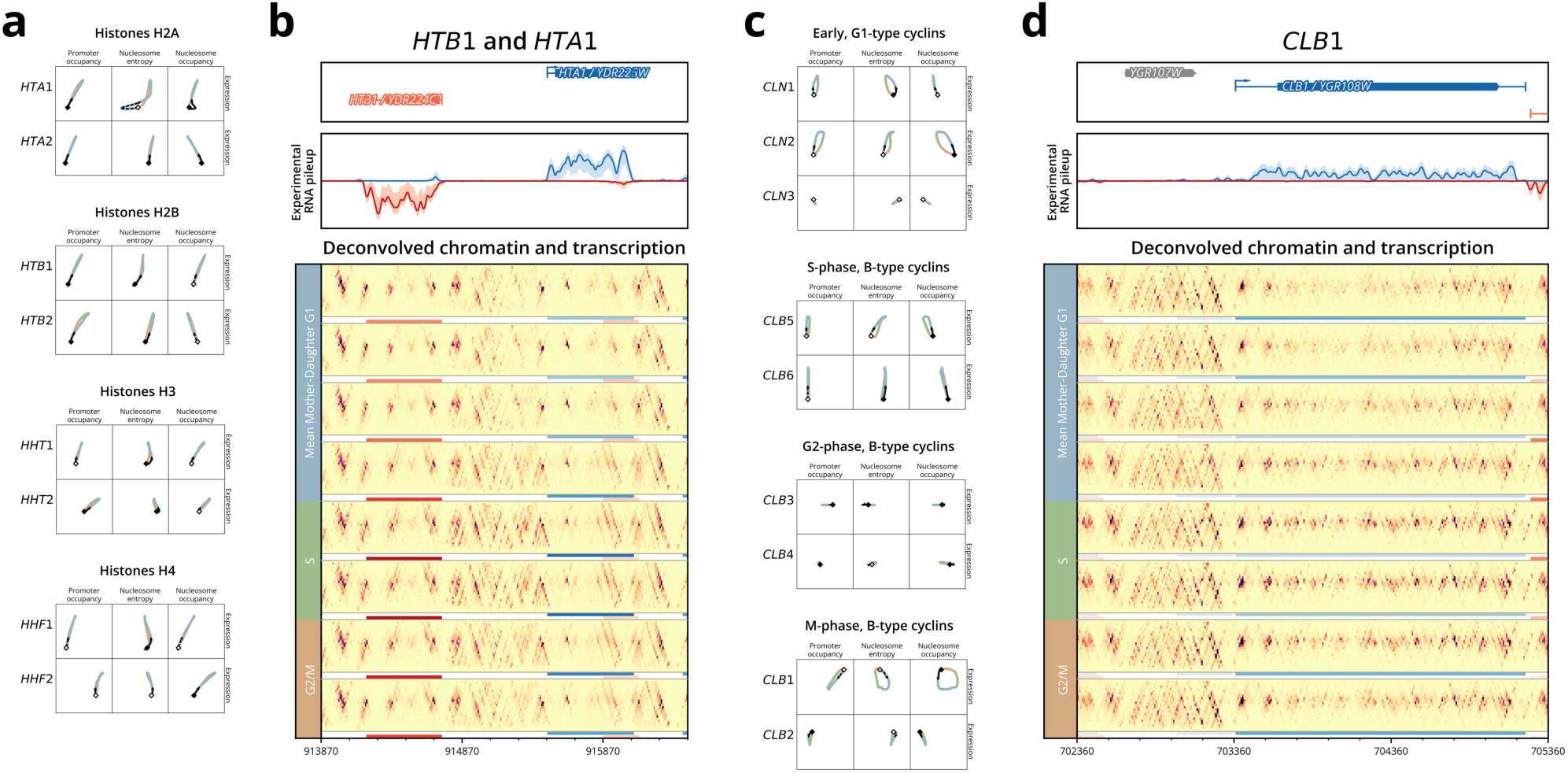
Cell cycle gene families exhibit diverse chromatin–expression coordination patterns. **(a)** Histone gene family trajectory plots show uniformly tight coordination between chromatin measures and expression. **(b)** Cyclin gene families display diverse regulatory patterns, exhibiting heterogeneous chromatin–expression relationships within subfamilies. **(c)** Locus-level view of histone genes *HTB1* and *HTA1* shows clear, characteristic chromatin dynamics for the histone gene family. **(d)** Detailed view of *CLB1* demonstrates clear nucleosome occupancy changes, with greatest changes occurring during S phase when expression is lowest.

Cyclins drive the cell cycle in four distinct activation waves (G1, S, G2, and M) (Barberis 2021; Bloom and Cross 2007). These cyclins interact with cyclin-dependent kinase Cdc28 to ensure tightly controlled cell cycle progression. For the genes within each wave, we observed generally coordinated chromatin-expression patterns (Fig. 6c). The S-phase cyclins are highly cycling in expression with moderate promoter changes, while the G2 cyclins are relatively unchanging. The most noticeable exceptions are within the Early G1 (G1-type) cyclins and M-phase (B-type) cyclins.

Compared to the other G1 cyclins which show clear cycling expression, *CLN3* shows unchanging chromatin and expression. This difference aligns with *CLN3*’s unique function as activator for *CLN1* and *CLN2* (Tyers *et al*. 1993). This dual role in promoting G1–S transition and in activating transcription may explain the requirement for greater stability in *CLN3*’s transcript levels. Among the mitotic cyclins, *CLB1* and *CLB2* show contrasting regulatory patterns: *CLB1* displays a clear lag between nucleosome dynamics and expression (Fig. 6c: *CLB1* right column), while *CLB2* cycles coordinately. This nucleosome lagging in *CLB1* indicates complex competing regulatory dynamics rather than simple coordinated regulation. Both genes are responsible for driving mitotic progression, but *CLB1*’s uncoupled nucleosome-expression relationship suggests more complex regulatory mechanisms are involved in its cell cycle expression (Fig. 6c,d).

Further, the MCM complex genes show heterogeneous regulatory patterns. In late G1, MCM proteins are loaded onto replication origins as part of pre-replication complex (pre-RC) formation, licensing these sites for DNA synthesis. The MCM complex then acts as the replicative helicase that unwinds DNA ahead of replication forks during S phase (Costa and Diffley 2022). While MCM genes share promoter elements and coordinate timing during G1/S (Hatoyama and Kanemaki 2023; Ohtani *et al*. 1999), our framework details substantial regulatory differences across complex members. These differences range from subtle, more stable changes, as with *MCM2*, to more significantly changing genes like *MCM7* (Supplemental Figure S10c), representing a particularly sophisticated example of cell cycle regulation.

During S phase, we observe subnucleosomal binding at a known Mcm1 site, identified by Rossi *et al*. (2021), precisely when *MCM7* expression is minimal. This observation is consistent with a well-characterized Mcm1 regulatory mechanism where Yox1 and Yhp1 bind with Mcm1 during S phase, forming a repressive complex that switches Mcm1 from an activator to a repressor (Pramila *et al*. 2002). Our chromatin deconvolution reveals the temporal dynamics of this repressive complex at the *MCM7* promoter.

Resolving this coordination—along with patterns across histones, cyclins, and other MCM complex genes— demonstrates CyCLOPS’s capability to systematically link chromatin changes to transcriptional regulation across diverse cell cycle pathways.

### Transcription factor binding cycles independently of target gene expression

While we demonstrated CyCLOPS’s capabilities in gene-level analysis, we were also interested in whether CyCLOPS could discern small-scale chromatin dynamics, such as transcription factor binding. Rossi *et al*. (2021) used a high-resolution ChIP-exo/seq assay to profile the genomic binding sites of approximately 400 different proteins in yeast. Of these proteins, we analyzed 61 factors with known transcriptional regulatory activity and DNA binding specificity, including 11 cell cycle TFs from Kelliher *et al*. (2018) (Supplemental Figure S11a). At these binding sites, we classified 685 of the 2,793 sites (25%) as cycling. We defined a threshold as the 95th percentile of all gene promoter binding occupancy PTR.

Cell cycle TFs showed no enrichment for cyclic binding compared to other TFs in the dataset, likely reflecting alternative regulatory mechanisms such as cooperative/competitive binding (Martin *et al*. 2023; Zhang *et al*. 2021) or post-translational modifications like phosphorylation (Whitmarsh and Davis 2000).

Beyond canonical cell cycle factors, several transcription factors exhibited unexpected cyclic binding patterns. These factors are each linked to the yeast metabolic cycle (YMC) and include Cha4 (serine/threonine metabolism) (Bonander *et al*. 2008), Hap5 (respiratory regulation, component of YMC) (Buschlen *et al*. 2003), and Urc2 (pyrimidine metabolism) (Andersen *et al*. 2008). The cyclic binding of these factors suggests a direct mechanism for coordinating YMC dynamics with cell division cycle progression.

Since transcription factors canonically regulate gene expression through promoter binding (Hahn and Young 2011), we asked whether cycling binding sites were enriched at promoters relative to all binding sites (Supplemental Figure S11b,c). Cycling TF binding sites showed promoter enrichment compared to the overall distribution (24% vs. 18%; Fisher’s exact p=0.014). Recent work by Mahendrawada *et al*. (2025) supports our findings that functional TF binding is concentrated at promoters. Despite this broad enrichment, only the pioneer factor Abf1 showed significant enrichment (p*<*0.005) for cyclic binding at promoters. These binding sites likely represent the functionally relevant binding events driving Abf1’s known role in mitotic cell cycle progression (Schlecht *et al*. 2008).

Having identified cyclic TF binding sites and their promoter enrichment, we asked whether cyclic TF binding correlates with cell cycle regulated gene expression. We examined 709 TF sites with binding within gene promoters. In specific cases, cell cycle regulated genes (Supplemental Figure S12a) show evidence for direct binding regulation, including the replication genes *CDC6* and *MCM7*, and histone genes *HHT1* and *HHF2* (Supplemental Figure S12d–f). However, genome-wide, cyclic binding does not directly correlate with cell cycle regulated expression (Supplemental Figure S12). This disconnect reflects the complexity of expression regulation, which may involve combinatorial TF control, chromatin priming, mRNA stability, and post-translational mechanisms beyond direct binding. Together, these findings reinforce that transcription factor binding and gene expression cycle largely independently during the cell cycle.

### Cyclic +1 nucleosomes link to transcriptional regulation with specialized histone modification signatures

To understand the dynamic regulation of nucleosomes, we examined 5,542 precisely identified +1 nucleosome positions from Chereji *et al*. (2018) (Supplemental Figure S13a). We computed three measures of nucleosome dynamics: entropy (disorganization), occupancy (presence/absence), and position (shifting), identifying the most stable and the most highly cycling 534 nucleosomes (10th and 90th percentile) (Supplemental Figure S13b).

Of the three measures of nucleosome cyclicity, we found a notable link between position cyclicity and cell cycle gene expression (Fig. 7a, Supplemental Figure S13c). Across position cyclicity deciles, the maximum expression PTR progressively decreased from the most dynamic to most stable nucleosomes. To understand this progression, we explored the role of the +1 nucleosome. The +1 nucleosome is well-understood as a regulator of gene expression. Its position establishes the nucleosome-depleted region (NDR) at promoters for TF and Pol II binding (Chereji and Clark 2018). While previous work did not detect this position-expression link (Li *et al*. 2026), CyCLOPS’s deconvolution reveals it by jointly modeling replicates with sub-minute resolution and copy number correction.

**Figure 7.**
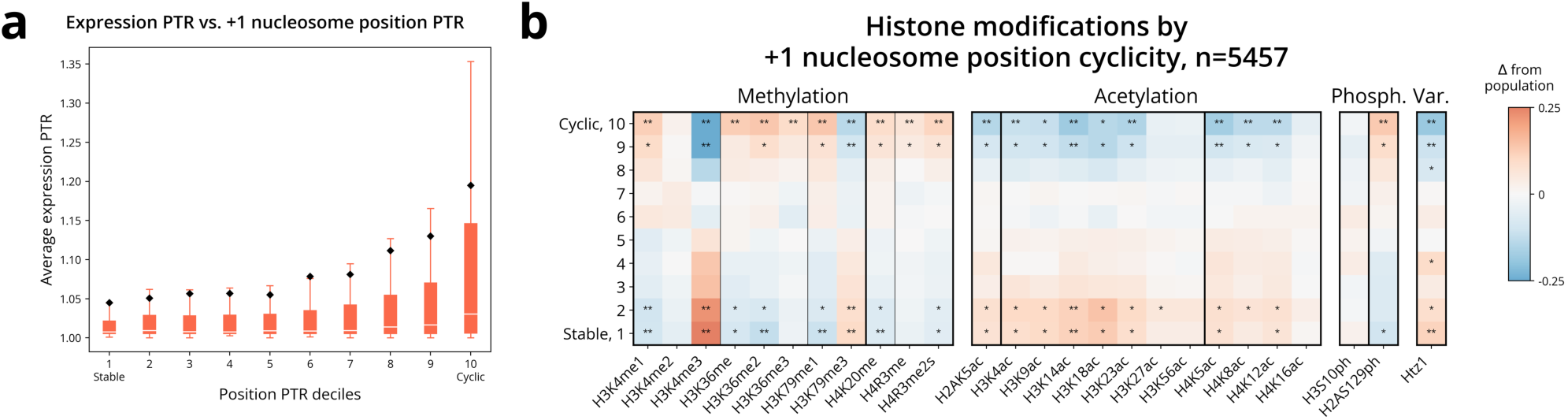
CyCLOPS reveals transcription-associated chromatin dynamics at cyclic +1 nucleosomes. **(a)** Distributions of expression PTR values progressively follow +1 nucleosome position cyclicity. Box plots of expression PTR values are plotted for each decile of +1 nucleosome position PTR. The features of the box plots progressively increase, moving from the most stably positioned to the most cyclically positioned nucleosomes. **(b)** Average histone modifications are also linked to position cyclicity. Modifications are plotted as a heatmap within deciles of position cyclicity. Among the most cyclically positioned nucleosomes, cyclic +1 nucleosomes show enriched H3K36 methylation and depleted acetylation, consistent with H3K36 methylation-mediated deacetylation that suppresses cryptic transcription (Butler and Dent 2012). For cyclically expressed genes, this mechanism prevents transcription from occurring during their inactive phases. Additionally, enrichment of H3K4me1 and H3K79me1 over their trimethylated forms reflects the incomplete methylation progression characteristic of genes that cycle between active and inactive transcription (Chory *et al*. 2019).

To explore the regulatory mechanisms of this positioning pattern, we examined histone modification markers from Weiner *et al*. (2015) across cyclicity deciles for each measure (Fig. 7b, Supplemental Figure S13d). While these measures reflect average histone modification levels in asynchronous cells, we found support for two established transcription-related mechanisms among position cycling nucleosomes.

First, H3K36 methylation-mediated deacetylation prevents cryptic transcription. During active transcription, Pol II-attached Set2p deposits methyl groups onto histone H3K36 (Butler and Dent 2012). This H3K36 methylation recruits the deacetylase complex Rpd3S to deacetylate nucleosomes and prevent cryptic, aberrant transcription. We observed these events reflected in enriched H3K36 methylation and depleted acetylation across cyclic +1 nucleosomes.

Second, for H3K4 and H3K79, enrichment of monomethylation over trimethylation is characteristic of genes with periodic expression patterns. Methyltransferases recruited to active promoters progressively methylate nucleosome histones, from unmethylated to mono-, di-, and trimethylated (me1/me2/me3 respectively), during transcription. For periodically expressed cell cycle genes, which undergo periods of active and inactive transcription, methylation may not progress to trimethylation due to either active demethylation or rapid nucleosome turnover (Chory *et al*. 2019). Consistent with this, we observe that cyclic +1 nucleosomes are enriched in H3K4me1 and H3K79me1 but depleted in their trimethylated forms.

Together, these results show that +1 nucleosome position dynamics are linked to cell cycle gene expression. The associated histone modifications confirm these position dynamics reflect real transcriptional regulatory mechanisms. CyCLOPS’s deconvolution revealed these previously hidden patterns.

### Non-genic cell cycle transcription exhibits links to subtle chromatin dynamics

In contrast to gene transcription, non-genic transcription represents a poorly understood regulatory phenomenon in eukaryotic cells. Changes in the chromatin may provide a window into understanding the mechanisms regulating these non-annotated transcripts. We next applied our framework to systematically analyze non-genic cell cycling transcripts and characterized the chromatin dynamics associated with their transcriptional changes.

We identified 256 non-genic transcripts spanning diverse genomic contexts, categorizing transcripts as antisense (transcribed on the opposing strand of a gene), divergent (sharing a promoter with an upstream opposing strand gene), both (exhibiting antisense and divergent contexts), or neither (Supplemental Figure S14a). Nearly half (109, 43%) occur without clear gene associations (“Neither” group), while antisense transcripts represent the next largest group (68, 27%), highlighting prevalence of antisense transcription.

To determine if these transcripts were cell cycling, we applied the same 95-percentile PTR threshold from the gene expression analysis. While representing 17% (44/256) of the total transcripts, the “Both” group makes up 38% (9/24) of cell cycling transcripts, suggesting non-genic transcripts considered both antisense and divergent interact with gene transcription in a synergistic manner during the cell cycle.

We were next interested in whether chromatin changes were linked to non-genic cell cycle transcription. By examining the chromatin PTRs for these transcripts, we again observed a poor linkage between chromatin changes and non-genic cell cycle transcription (Supplemental Figure S14a), mirroring the poor linkage we observed in gene expression. The stochastic nature of these non-genic transcripts likely provides an additional difficulty in systematically generalizing their behavior. However, by focusing on individual loci, we reveal a few clear examples of chromatin dynamics linked to non-genic transcription.

We found a divergent transcript sharing a promoter with S-phase gene *MCD1* (Supplemental Figure S14b). Both the divergent transcript and *MCD1* showed coordinated expression during late G1/S phase, consistent with *MCD1*’s expected cell cycle timing. During this period, +1 nucleosomes lost occupancy for both transcripts, while significant chromatin reorganization occurred throughout the shared promoter region. This coordination demonstrates how shared gene promoters can regulate non-genic transcription in the cell cycle.

This regulatory mechanism becomes more sophisticated in a transcript representing the “Both” category, antisense to *FPY1* and divergently transcribed from *SPT21* (Supplemental Figure S14c), genes both implicated during replication stress. This transcript benefits from two complementary chromatin effects. First, the stably transcribed *FPY1* likely provides consistent nucleosome displacement that enables antisense transcriptional machinery access. Second, the transcript shares a promoter with the cell cycle regulated *SPT21* gene. Moreover, this example shows the sequential timing of shared promoter activity. The non-genic transcript peaks in S phase after *SPT21* activation, suggesting it responds to shared promoter accessibility.

To determine the prevalence of this shared promoter phenomenon, we analyzed 75 divergent genenongenic transcript pairs, comparing cycling strength (using PTR) for each (Supplemental Figure S15a). We categorized the pairs into four groups based on the degree of concordant cycling, using a 95% PTR threshold. Only 3 pairs (4%) cycled together: the previously shown *MCD1* and *SPT21*, plus cell wall gene *SVS1* (Supplemental Figure S15b). Most other pairs (64/75, 85%) showed little to no cycling. For independently cycling pairs, we saw a modest enrichment for non-genic-only cycling (7 pairs) over gene-only cycling (1 pair). For example, cell wall remodeling gene *NCW2* maintains stable expression while an upstream non-genic transcript cycles (Supplemental Figure S15c). This enrichment for cycling non-genic transcripts suggests that shared promoters preferentially drive non-genic transcription.

Using CyCLOPS, we reveal diverse non-genic regulatory mechanisms—from promoter-specific control at divergently transcribed promoters to chromatin effects at antisense sites—that demonstrate how chromatin dynamics shape transcription beyond annotated genes.

### Deconvolution framework detects mother/daughter-specific chromatin differences

Asymmetric cell division in budding yeast requires daughter-specific processes such as cell wall degradation and bud site selection. Previous studies have systematically identified genes with daughter-specific expression patterns (Colman-Lerner *et al*. 2001). While this asymmetric expression is well described, we sought to examine whether the chromatin structure of these genes also differs between mothers and daughters.

While most of our analyses utilize averaged signals between mother and daughter branches, CyCLOPS models these branches separately and can discern their differences. To prevent misidentification of these differences, we applied stringent regularization to capture only the most significant mother/daughter asymmetries while filtering out technical variation.

We evaluated CyCLOPS’s ability to detect mother/daughter asymmetric patterns by examining daughter-specific genes. Interestingly, most daughter-specific expression genes exhibited subtle or no mother/daughter chromatin differences. Genes such as *DSE2* and *AMN1* showed minimal chromatin asymmetry despite their daughter-specific expression (Supplemental Figure S17b,d). The predominance of little to no chromatin differences suggests that regulating daughter-specific expression may occur primarily through mechanisms beyond chromatin-level control. This pattern also aligns with our previous observation that cell cycle chromatin dynamics and transcription changes act largely independently.

However, we identified a few genes with notable asymmetric chromatin patterns, including *SCW11* and *DSE1* (Supplemental Figure S17a,c). For *SCW11*, mother cells exhibited a greater nucleosome occupancy at nucleosomes flanking the TSS during G1 (Supplemental Figure S17f). The precise locations of these occupancy differences suggests a specific regulatory mechanism that prevents overexpression in mother G1 cells and indicates asymmetric chromatin-mediated control of gene expression. For *DSE1*, the most pronounced mother/daughter difference occurred downstream at a nucleosome within the middle of the gene body (Supplemental Figure S17e). This pattern suggests a mother-specific regulatory function during transcriptional elongation for *DSE1*.

These examples highlight the framework’s nucleosome-level precision in capturing mother/daughter-specific chromatin dynamics. More broadly, this component demonstrates CyCLOPS’s versatility in dissecting complex regulatory processes across multiple biological contexts.

## Discussion

Using CyCLOPS, we reveal the chromatin landscape throughout the cell cycle at unprecedented temporal resolution, uncovering both canonical regulatory dynamics and subtle patterns across diverse genomic contexts. CyCLOPS augments existing deconvolution methods (Guo *et al*. 2013) to use high-resolution chromatin data and implement a novel copy number correction procedure. We deconvolved high-fidelity replication profiles from the MNase data alone, with high correlation to established timing data (r=0.87). These profiles were then used to correct for pervasive copy number effects. Without this correction, we show copy number artifacts create false cycling signals and mask true dynamics (Fig. 2e,f). More broadly, this suggests that studies without comprehensive deconvolution and copy correction may systematically misinterpret cell cycle chromatin patterns.

To demonstrate CyCLOPS’s ability to resolve regulatory dynamics, we examined dynamics across cell cycle regulation contexts, both validating known cell cycle biology and uncovering subtle potentially novel regulatory mechanisms. We examined known cell cycle gene families with enhanced temporal resolution. This increased resolution enables a trajectory analysis of chromatin-transcription dynamics that reveals either direct coordination or temporally offset difference between chromatin and transcriptional changes. Analysis of histone genes revealed uniform chromatin-transcription linkage patterns across the gene family. In contrast, within cyclin and *MCM2–7* genes, we observed complex leading/lagging patterns between chromatin and transcriptional dynamics. These varied patterns illustrate the complexity of interpreting chromatin-transcription linkages, even among well-characterized cell cycle gene families.

Beyond well-characterized cell cycle gene families, our framework detects more subtle chromatin signatures in non-coding regions and in regions with mother/daughter-specific differences. For a small set of non-coding transcripts, we detected notable chromatin changes linked to cell cycle transcription patterns. We also identified subtle, potentially regulatory, mother-daughter chromatin asymmetries at individual nucleosome positions. These findings demonstrate the framework’s potential for characterizing subtle chromatin dynamics across diverse genomic contexts.

While CyCLOPS demonstrates substantial methodological advances, a few key assumptions limit its interpretation. First, our model separates and excludes recovery G1 following *α*-factor release from most of our analyses. By excluding this recovery phase, we limit our interpretation of mother/daughter G1 dynamics to the second and third cell cycle of our experiment. Second, the branching model design is limited in distinguishing mother and daughter G1, creating identifiability issues in accurate read assignment. We address this by applying a stringent regularization. This enables identification of meaningful mother-daughter differences, but assumes mother and daughter cells behave nearly identically through G1.

The current framework uses MNase-seq on *S. cerevisiae* as the model system. While MNase-seq captures rich factor-agnostic chromatin occupancy dynamics, our deconvolution framework can be extended to any time-series cell cycle experiment, including ChIP-seq, ATAC-seq, or methods that capture genomic state. Furthermore, the methodology is not limited to yeast—the core modeling approach is applicable to any eukaryotic system. CyCLOPS enables high spatiotemporal resolution of cell cycle chromatin dynamics, with broad potential to reveal new insights across diverse chromatin regulatory contexts.

## Materials and methods

### Deconvolution framework for cell cycle chromatin analysis

CyCLOPS extends two prior computational frameworks for cell cycle deconvolution. Orlando *et al*. (2007) first developed CLOCCS as a branching process using distinct recovery, mother, and daughter branches. This model estimated cell cycle parameters using a Markov-chain Monte Carlo (MCMC) algorithm against experimental budding index data of *S. cerevisiae* and *S. pombe*. Orlando *et al*. (2009) then further adapted this model for integration with flow cytometry data.

Guo *et al*. (2013) extended this approach by developing a deconvolution framework for gene expression. Using CLOCCS-derived cell cycle parameters, they constructed a convolution kernel matrix that jointly models independently collected experimental replicates of gene expression. Through a convex optimization approach, they deconvolve these coarsely sampled replicate experiments into idealized average single-cell gene expression profiles.

Building upon Guo *et al*. (2013)’s deconvolution framework, we developed CyCLOPS (Cycling Chromatin Landscape Occupancy Profiling System) to deconvolve the chromatin landscape throughout the cell cycle. We developed CyCLOPS in two key ways. First, while Guo *et al*. (2013)’s framework deconvolved individual gene expression profiles, our approach scales to genome-wide chromatin across millions of genomic positions. For assays like paired-end MNase-seq, this includes an extra dimension for fragment length analysis (Fig. 1b). Second, we addressed a challenge unique to genomic assays: unlike cellular RNA measurements, genome-based chromatin data are complicated by nonuniform copy number doubling during DNA replication in S phase.

These additions are built into the overall deconvolution framework in the following stages: (1) estimation of cell cycle parameters using the branching model (Fig. 1a) and MCMC algorithm of CLOCCS; (2) replication timing estimation from genomic MNase-seq occupancy; and (3) copy-corrected chromatin deconvolution (Fig. 1c). This approach enables idealized average single-cell-resolution analysis of diverse chromatin features through the cell cycle, with replication effects separated through copy number correction.

### Experiment and data preparation

We performed time course experiments using *α*-factor synchronization with sampling every 10 minutes over 150 and 140 minutes following release. At each time point, we collected MNase-seq samples (for chromatin analysis), RNA-seq samples (for transcript quantification), and flow cytometry samples (for cell cycle parameter estimation) (Supplemental Figure S1) in parallel from two independent biological replicates. All experimental protocols followed the procedures described in Li *et al*. (2021, 2026).

### Cell cycle branching model

We designed CyCLOPS based on the cell cycle branching process in Guo *et al*. (2013)’s framework, but adapted it specifically for *α*-factor synchronization experiments. The original model, designed for elutriated cells, consists of three distinct branches: initial, mother, and daughter. The initial branch includes a recovery period followed by a G1 phase shared with mother cells. This design is appropriate for elutriated cells that become essentially identical to mother cells following the recovery period.

For *α*-factor synchronization, however, two key modifications were required due to the distinct biology of this synchronization method. First, *α*-factor arrest creates fundamentally different recovery dynamics compared to elutriation (Ko and Moore 1990), necessitating a distinct recovery G1 (RG1) that differs from both mother G1 (MG1) and daughter G1 (DG1) (Fig. 1a).

Second, after release from *α*-factor, we found in our flow cytometry data (Supplemental Figure S1) a subset of cells that never leave the initial recovery G1. We model these cells using a “halted” state representing the initial population of cells that never initiate replication and progression into S phase.

#### Cell cycle parameter estimation and convolution kernel construction

We applied the CLOCCS (Characterizing Loss of Cell Cycle Synchrony) model (Orlando *et al*. 2009) to flow cytometry data to characterize how cell populations are distributed through the cell cycle at each experimental time point. To ensure accurate cell cycle parameter estimation, we fit CLOCCS with an extended burn-in period of 100,000 iterations and 100,000 iterations of sampling for each replicate (Supplemental Figure S2).

An additional parameter estimation step involves determining the delay parameter *α* between cytokinesis and cell wall breakdown. Flow cytometry will misclassify post-cytokinesis G1 cells as single G2 cells when cell walls remain intact, requiring us to estimate the duration of this delay for accurate M-to-G1 transition timing. Following Guo *et al*. (2013), we determined optimal *α* values using daughter-specific gene expression patterns, with an additional constraint requiring high correlation between replicates to ensure model stability. Details are provided in Supplemental Methods 7.

Following Guo *et al*. (2013), we used the CLOCCS-derived cell cycle parameters to construct convolution kernels **H**_1_ and **H**_2_ for each replicate. Each convolution kernel represents the proportion of the total cell population distributed across the idealized cell cycle profile (columns) over the experimental time course (rows) (Fig. 1c:middle).

The combined convolution kernel **H** was formed by vertically concatenating **H**_1_ and **H**_2_. This **H** kernel is subsequently used for deconvolution of replication timing, chromatin occupancy, and gene expression.

### Replication profile deconvolution

During S phase, genome replication creates variable copy numbers across the genome, which confounds interpretation of chromatin dynamics during the cell cycle (Fig. 2a). To address this challenge, we developed a deconvolution framework to estimate genome-wide replication timing profiles.

We constructed a matrix **G***_r_* to represent the broad genomic signal of copy number variation. **G***_r_* represents genome-wide MNase-seq occupancy organized as 10 kb windows (strided by 2 kb), windows broad enough to resolve replication dynamics. Each row corresponds to experimental time points and each column represents a genomic window (Fig. 2b; **G***_r_* left-most matrix).

After normalization at each time point, this matrix reveals oscillating regions of increased (red) and decreased (blue) occupancy across both the genome and time course. This effect is the interaction between variable DNA copy number during replication and the normalization constraint of equal total reads per sample.

We modeled **G***_r_* as a convolution between the unknown replication timing matrix **F***_r_* and the cell cycle convolution kernel **H**. Our model minimizes the L2-norm of differences between the modeled and observed occupancy (Fig. 2b).

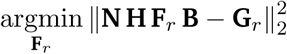

The replication model includes two essential terms: **N** corrects for the global normalization effect when total occupancy is held constant while actual DNA content increases during S phase (Supplemental Figure S3A:left). **B** represents the baseline occupancy level for each window, accounting for inherent differences in chromatin occupancy across the genome (Supplemental Figure S3A:bottom).

Given the stepwise nature and limited solution space of **F***_r_*, we used an exhaustive (brute force) search for the optimal profile. After optimizing **F***_r_*, the normalization matrix (**N**) and baseline occupancy matrix (**B**) were updated based on this optimization solution. We iteratively alternated the optimization of **F***_r_* and updates to **N** and **B** to ensure convergence to the final deconvolved replication profile **F***_r_*, representing the genome-wide replication timing across the cell cycle. Additional details for this alternating optimization procedure are provided in Supplemental Methods 3.

After applying a 2 kb stride to the genome, we obtained 6,045 genomic windows. We excluded 563 (9%) with insufficient read coverage (*<*75% threshold) and 112 (2%) lacking adequate signal for reliable timing estimation. In total, we estimated replication timing for 5,370 windows.

### Deconvolution of the chromatin

CyCLOPS applies the CLOCCS-derived convolution kernel to recover idealized average single-cell chromatin profiles from population-level measurements. The deconvolution model solves an optimization problem to identify the chromatin profile that best reproduces the observed experimental data when convolved with the population distribution, encoded in the convolution kernel **H** (Fig. 1c).

#### Chromatin deconvolution model

We modeled chromatin occupancy **G***_c_* as the convolution between the estimated cell cycle proportions per time point, encoded in the convolution kernel **H**, and the idealized average single-cell chromatin occupancy profiles **F***_c_* (Fig. 1c).

To enable simultaneous analysis of multiple chromatin occupancy values, we transformed the MNase-seq data into multidimensional histograms representing fragment length-specific chromatin occupancy across the genome. Our model extends the one-dimensional deconvolution approach of Guo *et al*. (2013) by augmenting the input and output vectors into two-dimensional matrices **G***_c_* and **F***_c_*, where the *m* columns act as independent measurements of chromatin occupancy.

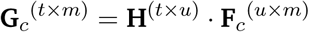

Our optimization is set up as an objective function that identifies the average single-cell profile **F***_c_* that minimizes the difference between the estimator **H** *·* **F***_c_* and observed read counts **G***_c_*. The *m* columns in **G***_c_* and **F***_c_* act as independent minimization objectives.

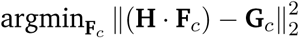

To ensure biologically plausible solutions, we regularized the objective function to enforce three biological constraints: (1) temporal smoothness of chromatin changes across cell cycle phases, (2) similarity between mother and daughter G1, and (3) similarity between halted cells and recovery G1 cells.

The updated, regularized objective function is defined as:

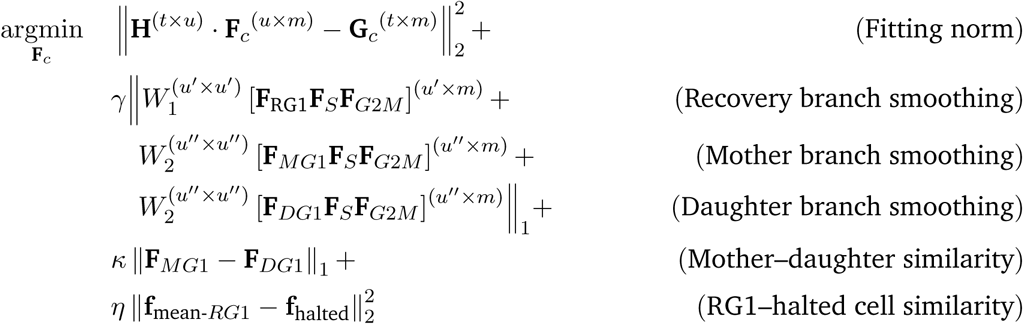

We enforced smoothness using wavelets as described in Guo *et al*. (2013). The wavelets transformed the solution onto a local frequency domain where applying an L1 penalty encouraged smoother fits. We used the same Symlet wavelets (with parameter *n* = 5) selected by Guo *et al*. (2013) for their smoothness and symmetry properties. We weighted this regularization term using parameter *ц*. The mother-daughter G1 constraint addresses a key identifiability challenge: while we aim to model the distinct daughter G1 separately from mother G1, the model may lack sufficient information to reliably distinguish between these two G1 states. Thus, we regularize this difference to ensure a high level of similarity between mother and daughter branches. This constrained flexibility allowed us to identify distinct mother/daughter-specific differences. We weighted this regularization term using the parameter *n*.

In our *α*-factor arrest experiment, we found that a proportion of cells never exited the initial recovery G1. These halted cells are evident in both the flow cytometry data (Supplemental Figure S1a,b), as the persistence of single copy DNA cells, and in the CLOCCS-parameter estimations (Supplemental Table 1), estimated as 23% and 29% halted cells. These halted cells are likely nearly identical to recovery G1 cells, but are unable to initiate replication. To enforce this similarity, we applied L2 regularization to constrain the differences between halted cells and the average of the cells in recovery G1, weighted by *η*.

Details for the selection of each of these regularization parameters are described in Supplemental Methods 5.

#### Incorporation of copy number correction

To distinguish true regulatory chromatin changes from replication artifacts, we incorporated the deconvolved replication profiles directly into our chromatin deconvolution model.

In this integrated approach, we model the observed chromatin occupancy data (**G***_c_*) as **NH**(**F***_c_* ⊙ **F***_r_*)**B**, where **F***_r_* scales the chromatin signal to account for DNA replication effects. Here ⊙ denotes element-wise multiplication. This scaling captures the expected doubling of DNA content in newly replicated regions as cells progress through S phase into G2/M, while preserving the underlying regulatory chromatin dynamics encoded in **F***_c_*.

The updated objective function separates copy number effects (encoded in **F***_r_*, **N**, and **B**) from non-replicative chromatin dynamics (**F***_c_*):

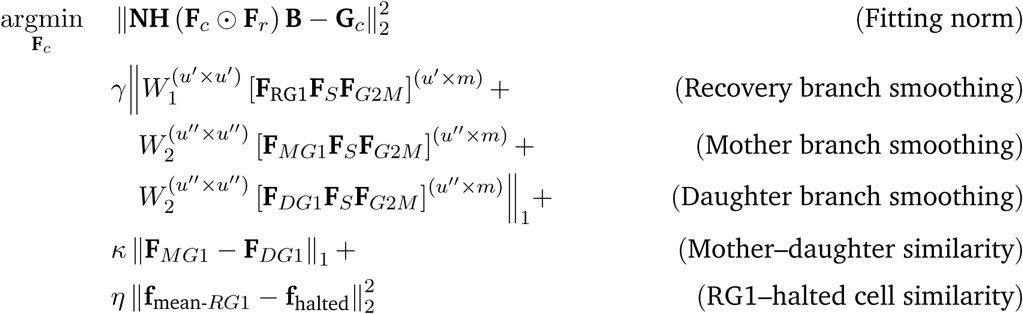

This copy number-corrected deconvolution ensures that observed chromatin changes reflect true chromatin dynamics rather than changes due to copy number effects. The resulting framework enables genome-wide recovery of average single-cell chromatin profiles from population-level measurements, providing the foundation for accurately characterizing chromatin dynamics throughout the cell cycle.

### Calculating measures of the chromatin for genes

We calculated three complementary metrics to characterize chromatin dynamics across gene regions. We defined two regions for analysis: promoter regions spanning 200 bp upstream of the transcription start site (TSS) (Supplemental Figure S9), and gene body regions extending 500 bp downstream of the TSS to encompass the first three nucleosomes.

First, we defined two measures of chromatin occupancy: small fragment promoter occupancy and gene body nucleosome occupancy. Small fragment promoter occupancy was defined as fragments of length 0–100 bp within promoter regions. These small fragments reflect transcription factor binding and other sub-nucleosomal DNA binding factors at gene promoters. Nucleosome occupancy was defined as reads 130–200 bp long within gene body regions. These reads represent nucleosome presence relating to DNA-dependent processes such as transcription and replication.

For nucleosome reads, we computed an additional measure, gene body nucleosome entropy, a measure of nucleosome position variability previously used by Tran *et al*. (2021):

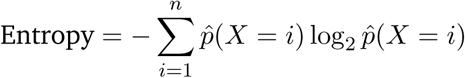

where *p*^(*X* = *i*) represents the probability of nucleosome occupancy at each position *i* within the region of size *n*. This models nucleosome positioning as a probability distribution and measures the uncertainty in nucleosome position. Higher entropy values indicate more variable positioning, while lower values reflect more organized nucleosome arrangements. Because the metric is normalized into a probability distribution, it is resistant to broad occupancy-related artifacts.

### Calculating peak-to-trough ratio (PTR) to measure strength of cycling

We calculated peak-to-trough ratio (PTR) to quantify the strength of cell cycle oscillations in both chromatin metrics and gene expression. PTR was computed as the ratio of the 90th percentile to the 10th percentile value to minimize the influence of outlier measurements.

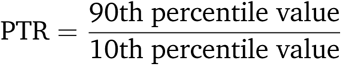

We calculated PTR for both experimental data (clock time) and deconvolved profiles (deconvolved time). For experimental data, we calculated PTR using measurements from the second and third cell cycles, excluding the recovery period (0–30 min for replicate 1 and 0–20 min for replicate 2) which exhibits *α*-factor-related effects. For deconvolved profiles, we calculated PTR using the average of the mother and daughter branch profiles, again excluding the *α*-factor effects on the recovery branch.

### Normalized trajectory area calculation

To quantify the correlation strength between chromatin dynamics and gene expression in cell cycling genes, we calculated normalized trajectory areas from paired time series measurements. For each gene, a chromatin measure and the gene’s expression values across the deconvolved cell cycle time course are considered as two-dimensional trajectories. The area of these trajectories measures the correlation strength between expression and the chromatin metric. Linear, small area shapes represent strong direct correlation, whereas circular, larger areas represent leading and lagging timing between the measures.

The area enclosed by each trajectory was calculated using the shoelace formula for polygon area (Braden 1986). Trajectory areas were normalized by the square of the maximum distance between any two points in the trajectory (diameter-squared normalization) to standardize the measure for ranking the strength of chromatin-expression coupling in cycling genes. This normalization reduces false positives generated from long elliptical areas and ensures that the larger area values correspond to circular shape trajectories.

### Transcript calling procedure for non-genic transcript identification and gene TSS refinement

We used stranded RNA-seq data to find non-genic transcripts and precisely refine gene TSS positions using a transcript calling approach. We defined transcript boundaries using a two-threshold approach applied to the stranded RNA-seq pileup data. Primary thresholds identified transcript seeds, and extension thresholds expanded these seeds to natural boundaries. Nearby regions with similar average expression levels were merged into called transcripts. We obtained final transcript calls if the length was 200 bp or longer, thereby excluding spurious short transcripts.

We identified 256 non-genic transcripts that did not overlap with any same-strand gene boundary and updated 3,507 gene TSS positions. These genes were updated by comparing RNA-seq transcript calls to existing TSS annotations.

Prior to our refinement procedure, we defined annotated TSS positions using Park *et al*. (2014) if available, and otherwise by 5’ ORF boundaries in sacCer3/R64 (Engel *et al*. 2022). In both cases, a gene TSS is updated if our RNA call was within 200 bp of the prior annotation (Supplemental Figure S16a). This 200 bp threshold captured the most well-defined transcript calls, validated by visual inspection of transcript and promoter architecture agreement (Supplemental Figure S16b,c).

## Code availability and data accession

Code for CyCLOPS is available at https://github.com/HarteminkLab/chromatin-deconvolution-paper. All genomic data are available at the NCBI GEO repository with the accession number GSE168699.

## Acknowledgements

We thank current and former members of the Hartemink group and the MacAlpine lab for critical comments and suggestions throughout the entirety of the project. T.Q.T., Y.L. and A.J.H. were supported by National Institutes of Health (NIH) grant R35-GM141795, and D.M.M. was supported by NIH grant R35-GM127062.

