## Supplemental Materials for "Deconvolving the spatiotemporal chromatin landscape through the cell cycle"

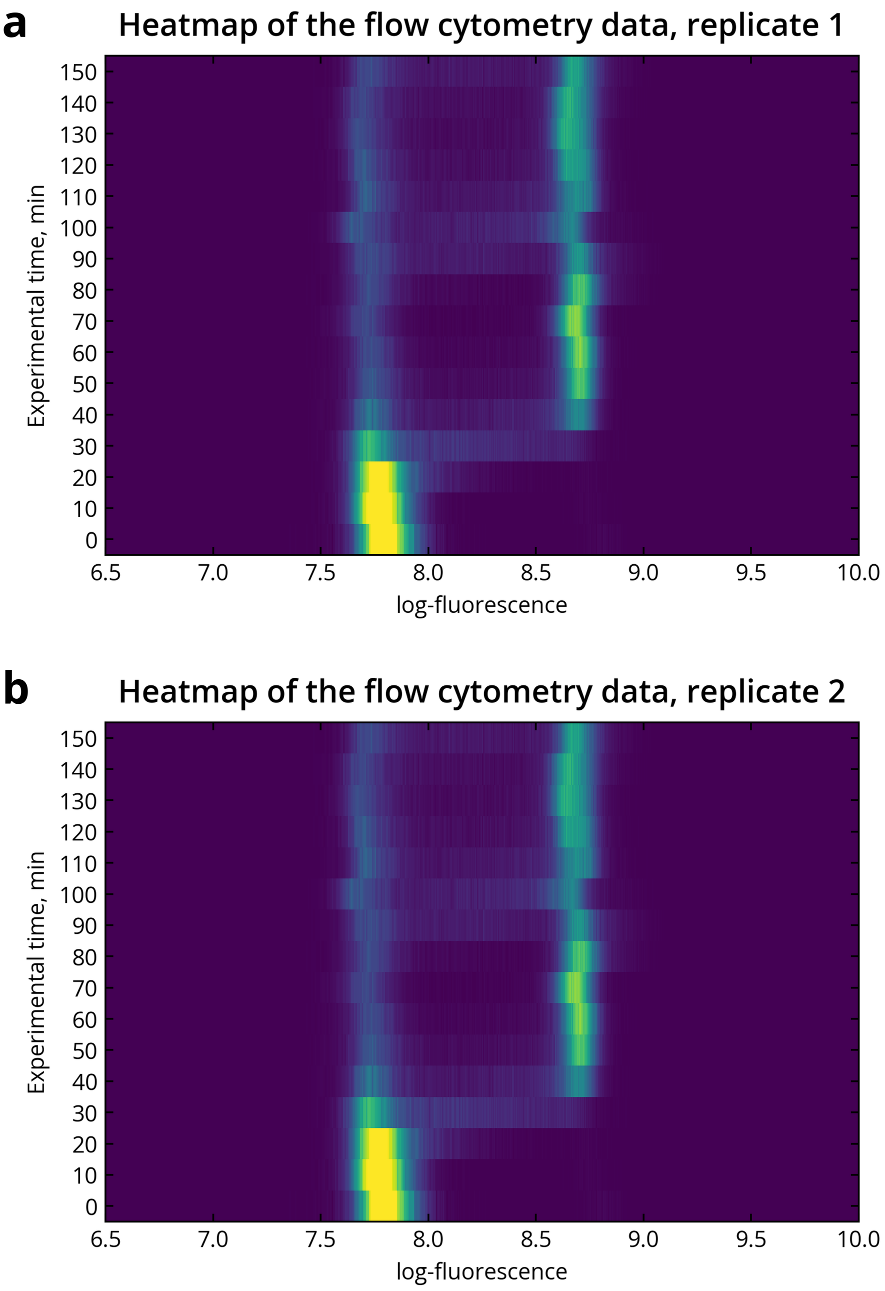


**Supplemental Figure S1**. Heatmaps depicting flow cytometry data for experimental replicates 1 **(a)** and 2 **(b)**. Heatmaps show cell cycle synchrony in DNA content throughout the time course. Log-fluorescence is plotted on the x-axis representing DNA content. Cells in G1 have log-fluorescence values near 7.7, while cells in G2 have double the DNA content with log-fluorescence values near 8.7. Each time course is sampled every 10 minutes up the y-axis for 150 minutes.

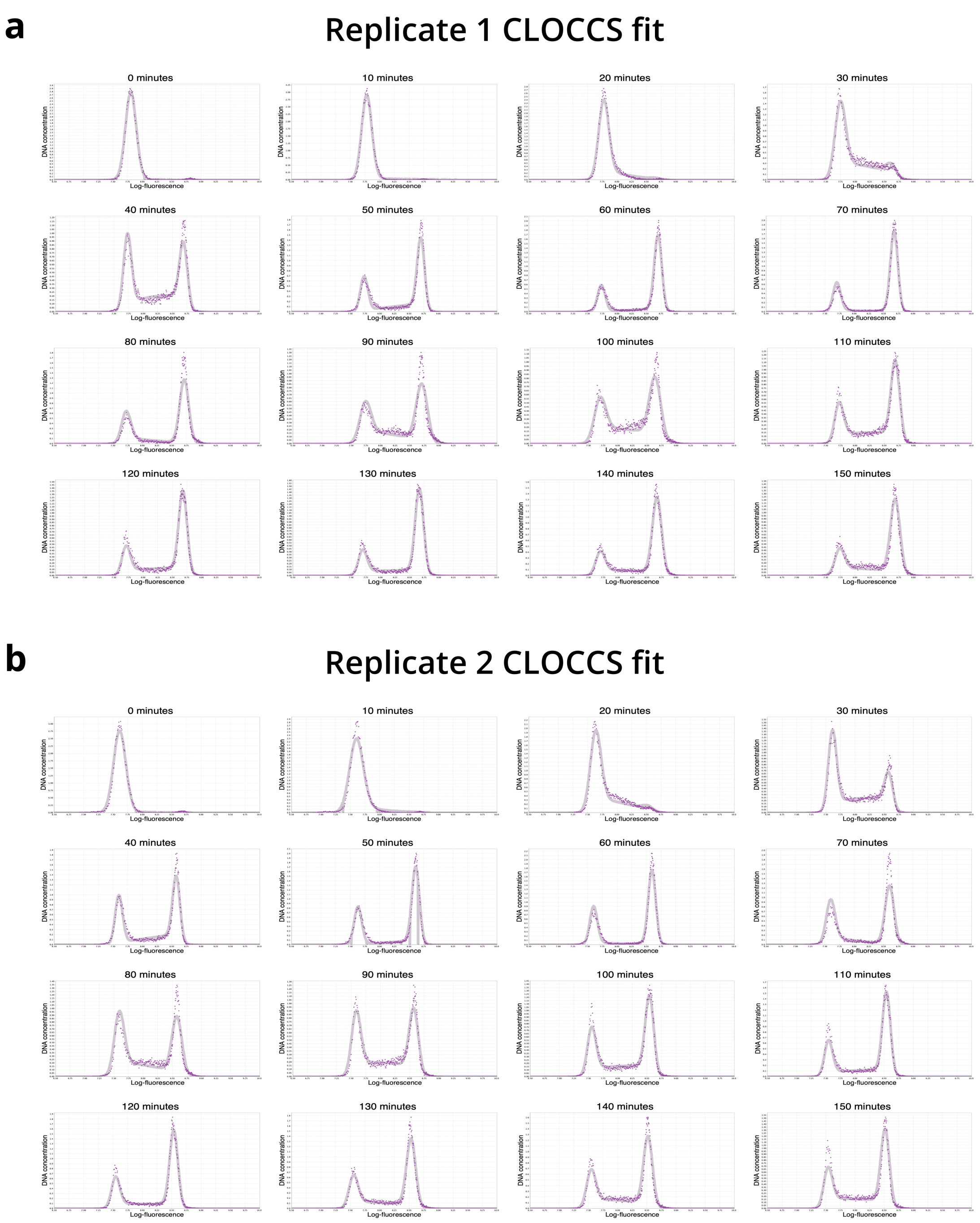


**Supplemental Figure S2**. Extended runs of cell cycle estimation model CLOCCS generate curve fits to flow cytometry data. These fits largely agree with the raw data in replicates 1 **(a)** and 2 **(b)**. At each time point, the raw data (purple points) are plotted atop the CLOCCS fit (gray curve). The 95% confidence interval of the fit is indicted by the thickness of the gray curve.

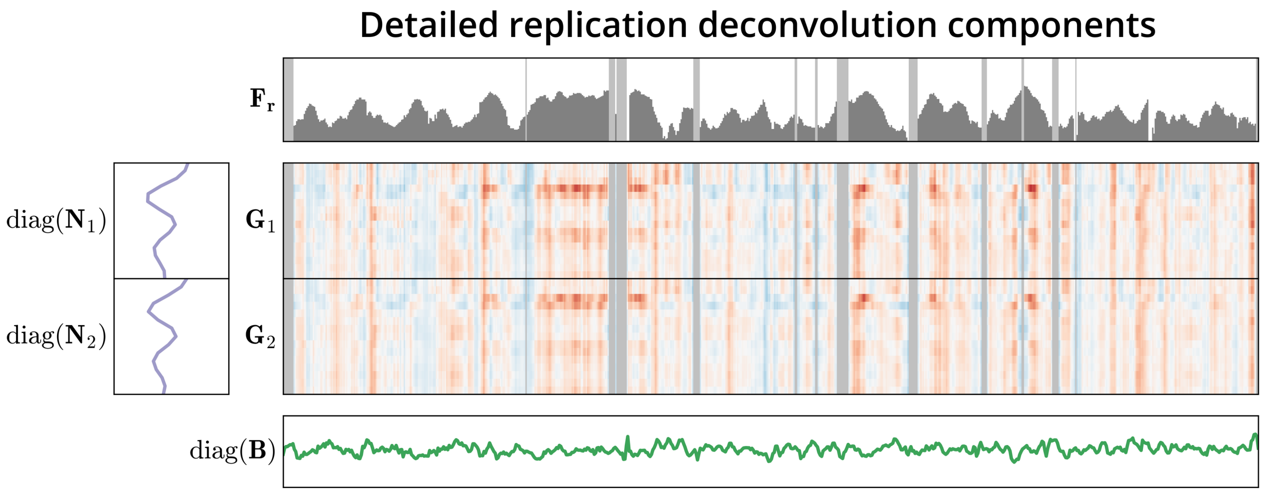


**Supplemental Figure S3**. Diagonal normalization matrices **N** and **B** provide additional context needed for replication deconvolution. Top: Deconvolved replication profile $\mathbf{F}_r$ shows the time (increasing downward) at which windows of the genome transition from one copy (white) to two copies (dark gray). Left: **N**, composed of $\mathbf{N}_1$ and $\mathbf{N}_2$, corrects for the average amount of replicated DNA at each time point for each replicate. Middle: Normalized MNase-seq occupancy **G**, composed of $\mathbf{G}_1$ and $\mathbf{G}_2$, encodes replication timing for each replicate. Regions with low-read density are masked (light gray). Bottom: Baseline occupancy **B** corrects for variation in per-window occupancy across the genome.

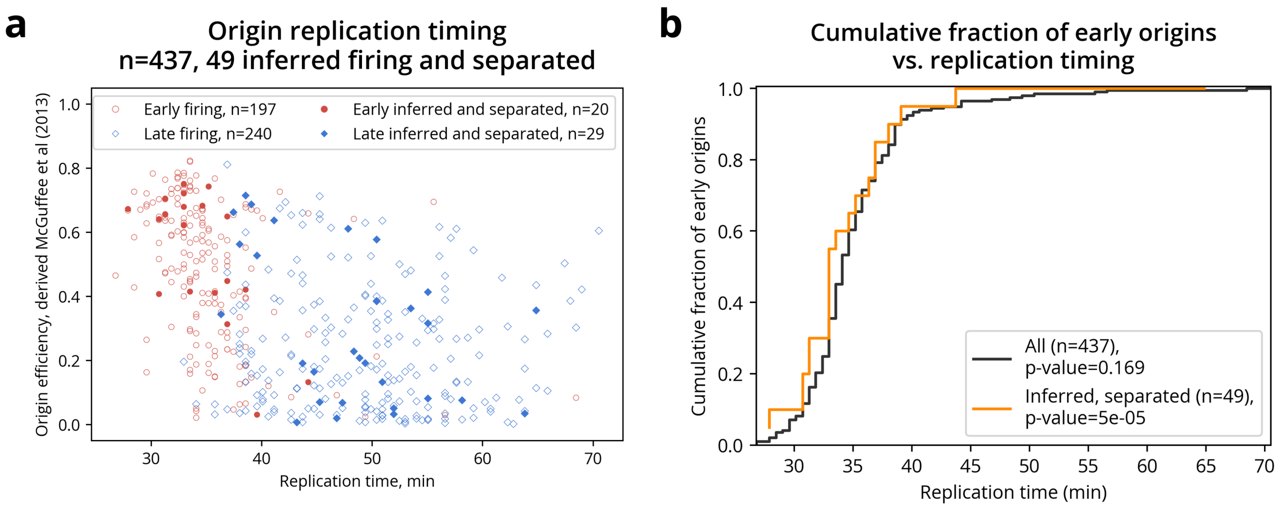


**Supplemental Figure S4**. Replication profile estimation agrees with known early and late origin annotations. **(a)** Origin replication timing correlates with annotated early and late origin classifications. Replication timing estimates (x-axis) are plotted against origin efficiency derived from McGuffee *et al.* (2013) (y-axis). Points are colored by early/late annotations from Belsky *et al.* (2015). Filled circles indicate "inferred firing" origins where local replication timing was earlier than genomic neighbors. **(b)** Cumulative fraction of early annotated origins across replication timing validates filtering approach. Following a permutation test for statistical significance, we found that early origins show insignificant enrichment among early replicating regions when analyzing all 437 annotated origins (black line, $p=0.169$). However, filtering to 49 inferred firing and separated origins (those replicating earlier than neighboring regions and $>5$ kb from other origins) reveals highly significant enrichment (orange line, $p<0.0001$).

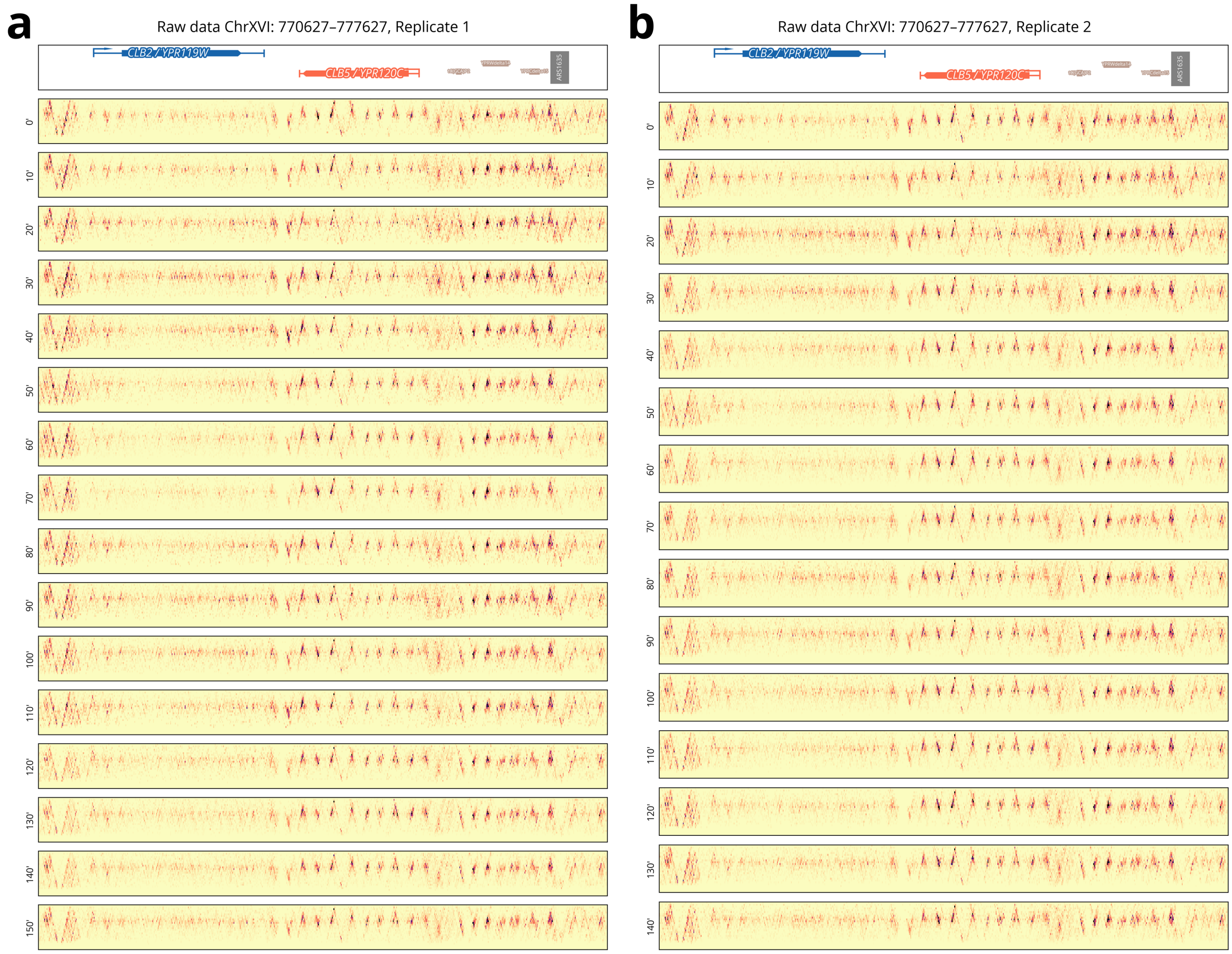


**Supplemental Figure S5**. Raw chromatin data for *CLB2* and *CLB5*. Time course data for replicates 1 **(a)** and 2 **(b)** show similar signals of cell cycle chromatin changes. For example, in both replicates, the gene body of *CLB2* shows a clear signal of cyclic nucleosome disruption and organization. CyCLOPS aims to capture consistent signal shared across both replicate experiments.

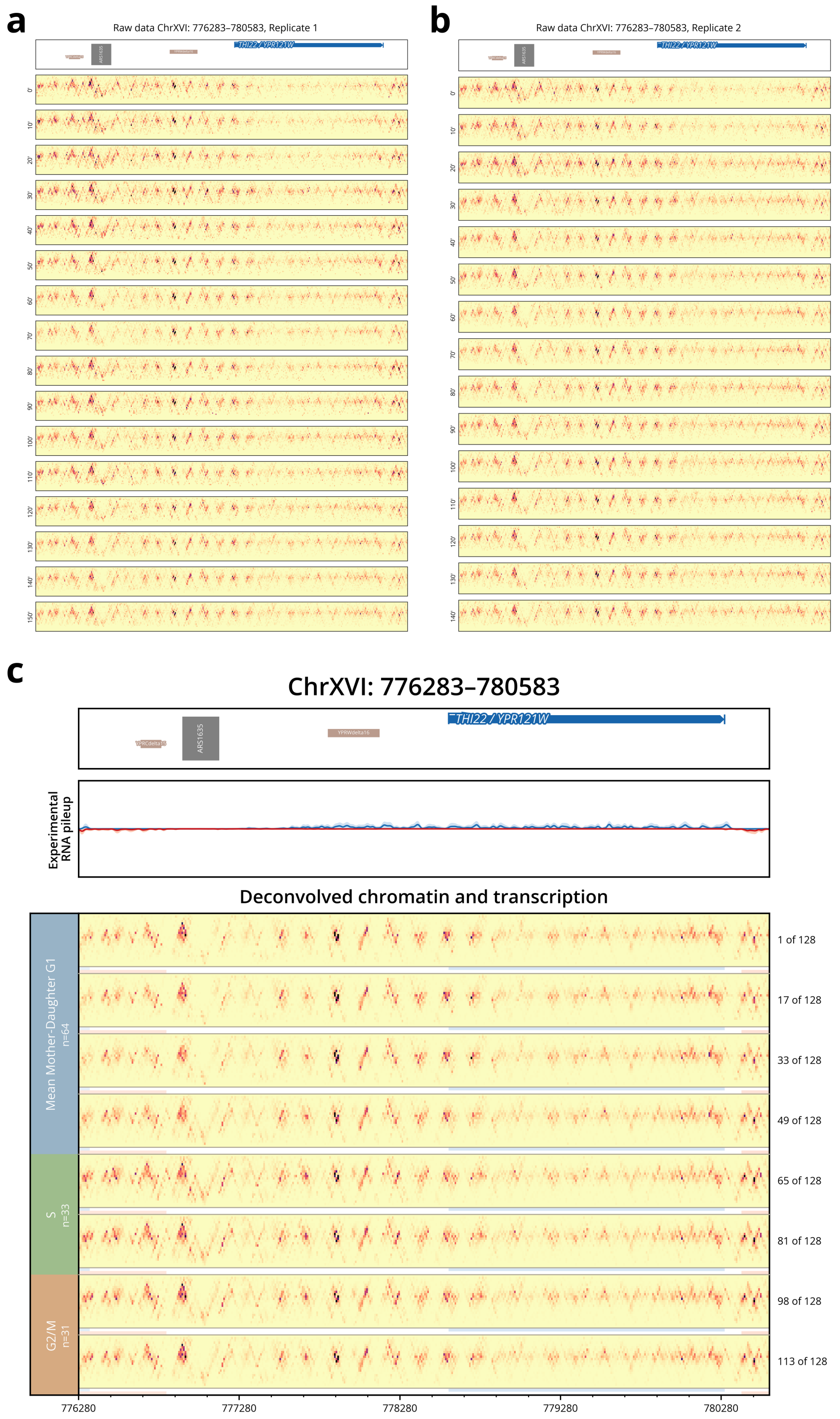


**Supplemental Figure S6**. **(a,b)** Raw chromatin data across two replicate experiments for *THI22*, a gene consistently expressed at a low level over the course of the cell cycle. **(c)** Deconvolved chromatin signal for *THI22* shows unchanging chromatin for this lowly transcribed gene. In contrast, upstream (to the left) of *THI22*, the origin *ARS1635* shows a clear signal of increased footprint occupancy prior to and during S phase.

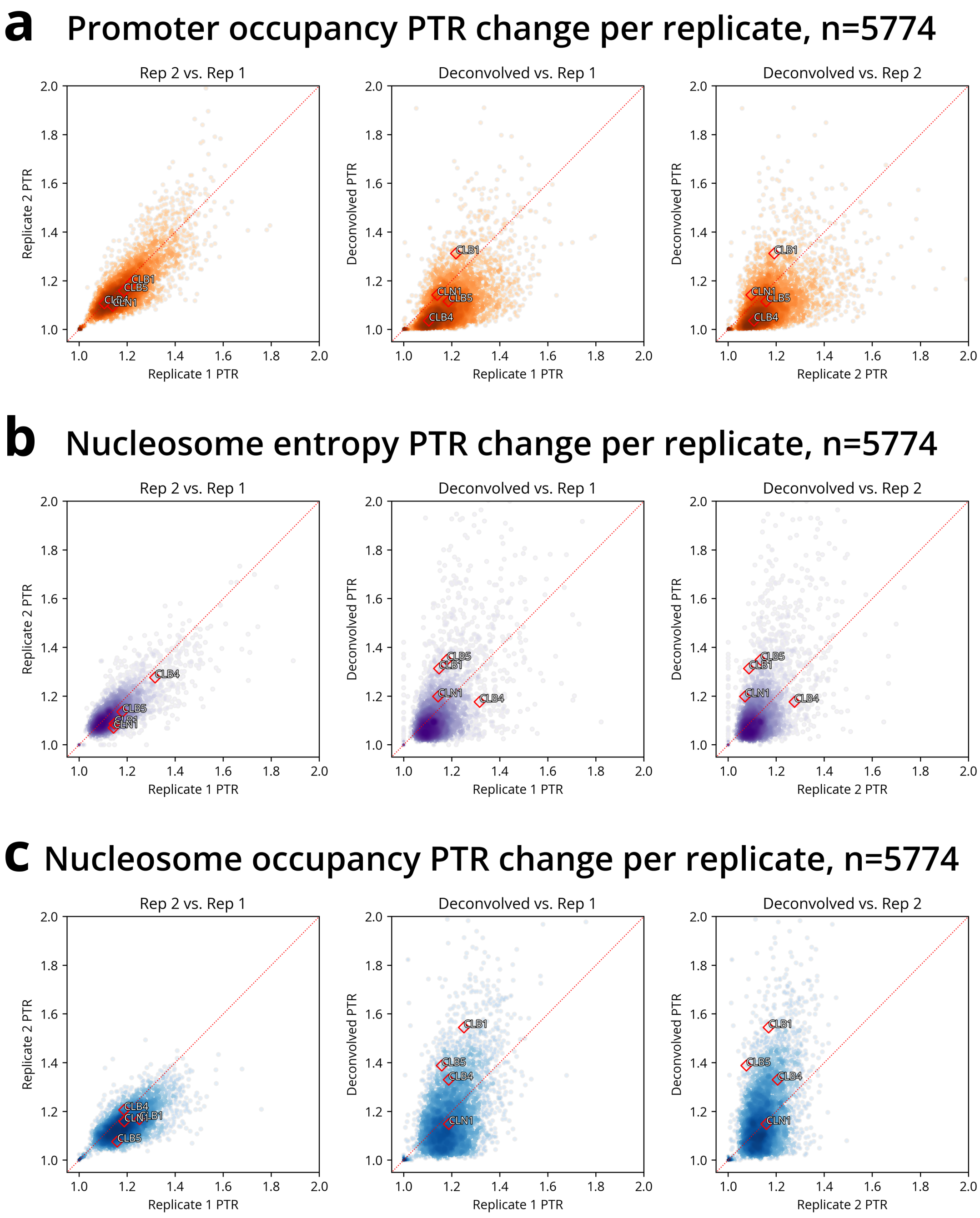


**Supplemental Figure S7**. Deconvolution reveals true cell cycle chromatin changes. Scatter plots compare cyclicity (PTR) across biological replicates for **(a)** promoter occupancy, **(b)** nucleosome entropy, and **(c)** nucleosome occupancy. Left panels show concordance of PTRs from replicate data before deconvolution. Middle and right panels compare deconvolved PTRs to each replicate's PTR. Cyclins *CLN1*, *CLB1*, *CLB4*, and *CLB5* with cyclic signal are plotted with red diamonds for reference. Deconvolution broadly increases PTR for nucleosome metrics (b, c: most points above diagonal), indicating improved cell cycle signal. In contrast, promoter occupancy shows mixed effects following deconvolution, with many genes decreasing in PTR. We found that small fragment signals were more susceptible to noise in the raw data, so the PTR reduction may reflect the removal of false cyclicity signal.

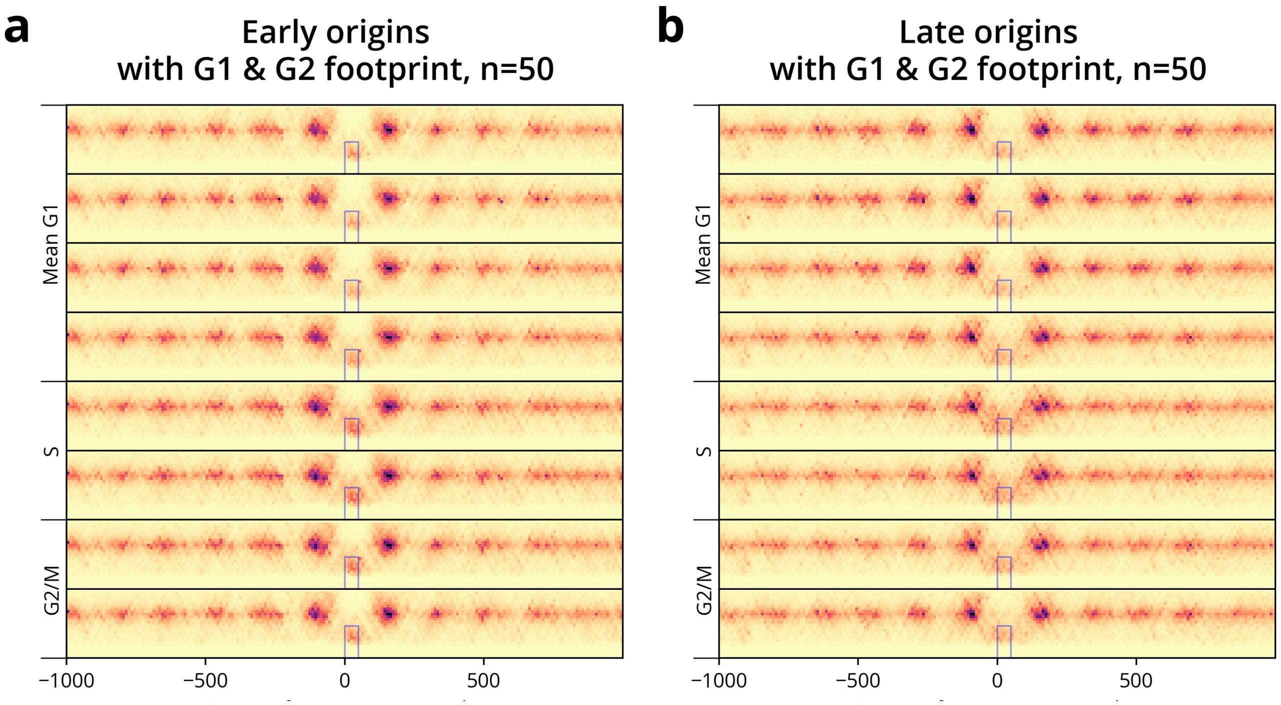


**Supplemental Figure S8**. Early firing origins show greater signal of pre-RC formation than late firing origins. **(a)** Composite profile is formed from the 50 earliest firing origins with G1 and G2 footprint classifications from Belsky *et al.* (2015). Early origins show a prominent small fragment footprint in S phase. **(b)** The equivalent composite for the 50 latest firing origins shows a similar but reduced footprint signal, consistent with lower firing efficiency relative to early origins.

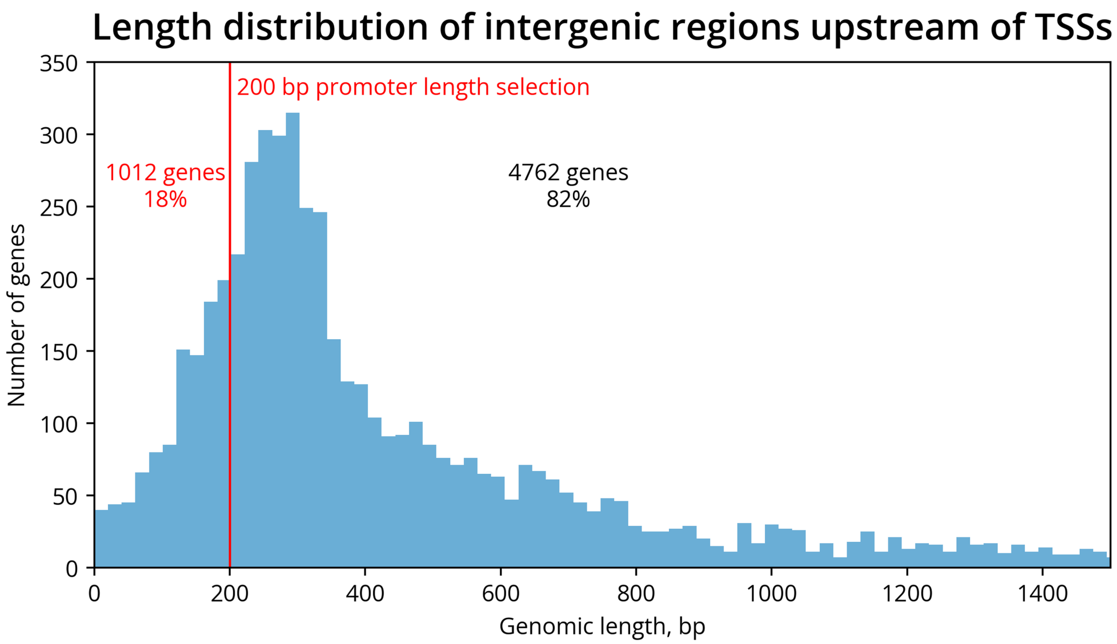


**Supplemental Figure S9**. Promoter length for analysis is selected based on intergenic spacing. For each gene, the distance to the nearest upstream gene (on either strand) was computed and plotted as a histogram. A 200 bp promoter region was selected to ensure that 82% of genes have sufficient intergenic space, avoiding overlap with upstream gene bodies.

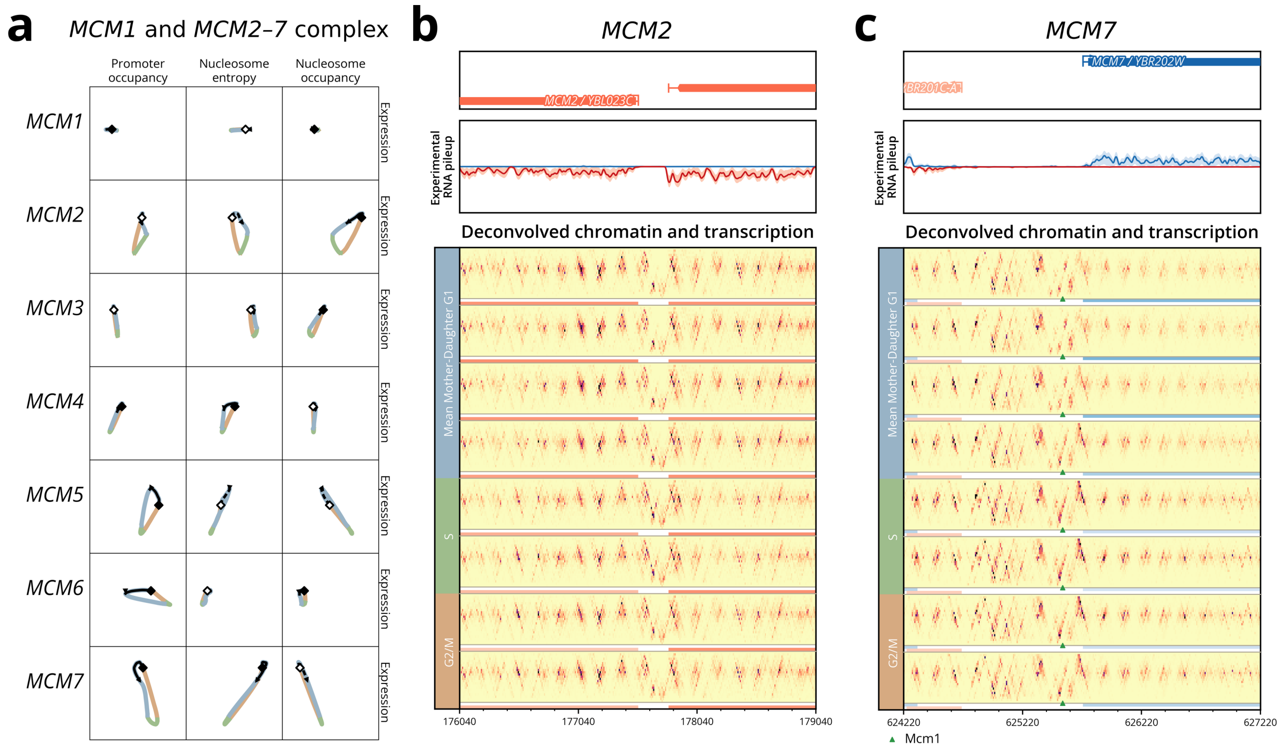


**Supplemental Figure S10**. The *MCM2–7* complex genes exhibit heterogeneous chromatin-transcription coordination. **(a)** Overview of all *MCM2–7* complex members, along with non-complex member and transcriptional regulator *MCM1*. Individual *MCM* genes display diverse regulatory patterns ranging from minimal to substantial chromatin dynamics. **(b)** *MCM2* displays relatively stable expression and subtle chromatin changes throughout the cell cycle. **(c)** *MCM7* demonstrates strong expression cycling with coordinated chromatin dynamics. Repressed expression in S phase coincides with promoter binding at a known *MCM1* binding site (green triangle), consistent with Mcm7/Mcm1-mediated autoregulation.

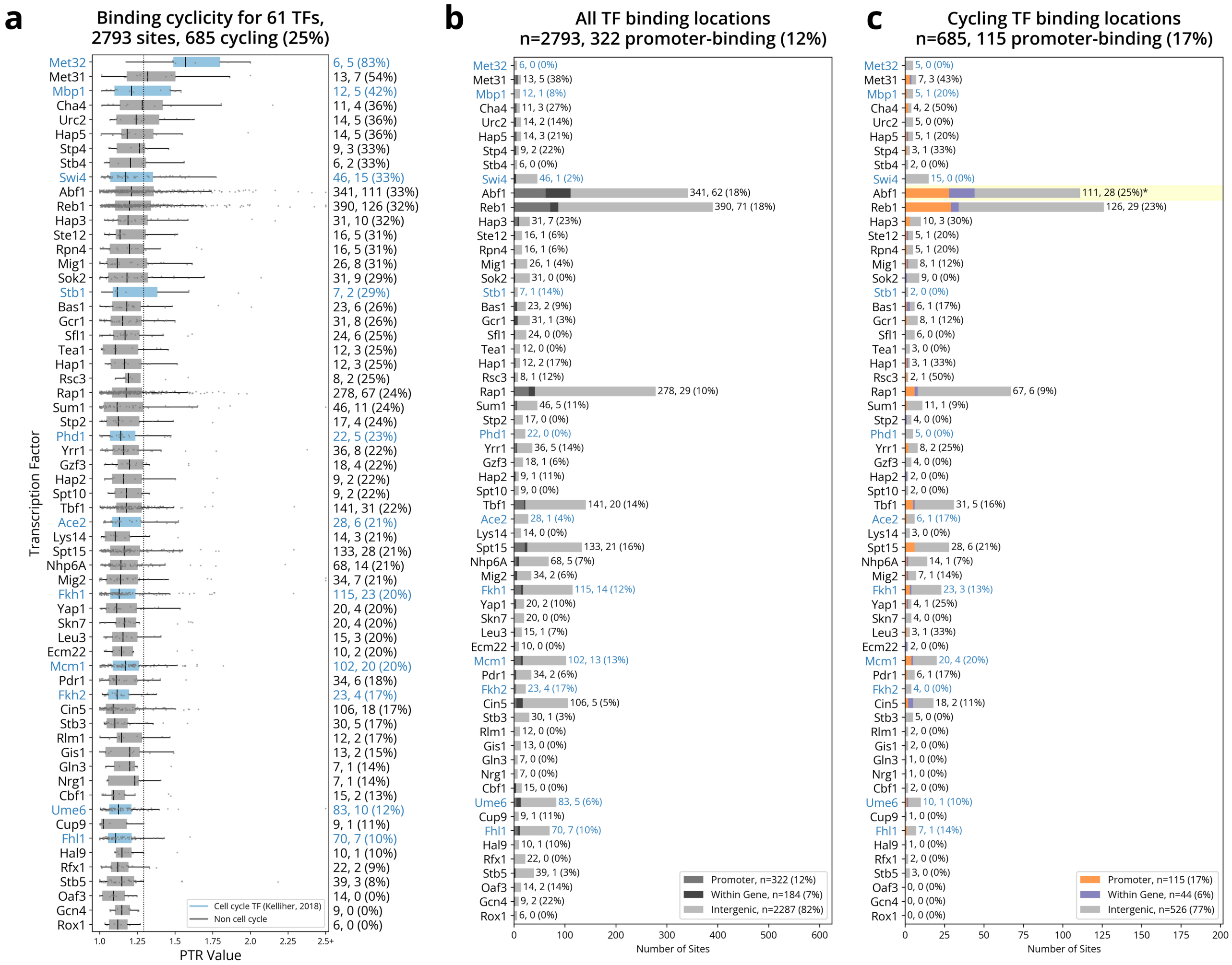


**Supplemental Figure S11**. Cell cycle transcription factors (TFs) do not exhibit more frequent binding cyclicity than other TFs. **(a)** Peak-to-trough ratio (PTR) analysis of 61 TFs across 2,793 binding sites from Rossi *et al.* (2021). 685 of these binding sites (25%) are classified as cycling, defined as exceeding the 95th percentile threshold from the distribution of promoter occupancy PTRs. TFs are ranked by their proportion of cyclic binding sites. Cell cycle TFs (shown in blue) from Kelliher *et al.* (2018) show no enrichment for cyclic binding compared to other TFs. Notably, metabolic cycle-associated factors (Cha4, Hap5, Urc2) exhibit unexpected cyclic binding patterns. **(b)** Distribution of binding site locations shows that a small proportion (12%) of TF binding occurs in gene promoters. **(c)** Binding sites that are cycling show a largely similar distribution across binding locations, with a slightly higher 17% binding in gene promoters. Among individual factors, only the pioneer factor Abf1 shows statistically significant enrichment ($p<0.005$), with 25% of its cyclic binding occurring in gene promoters, consistent with Abf1's role in mitotic progression.

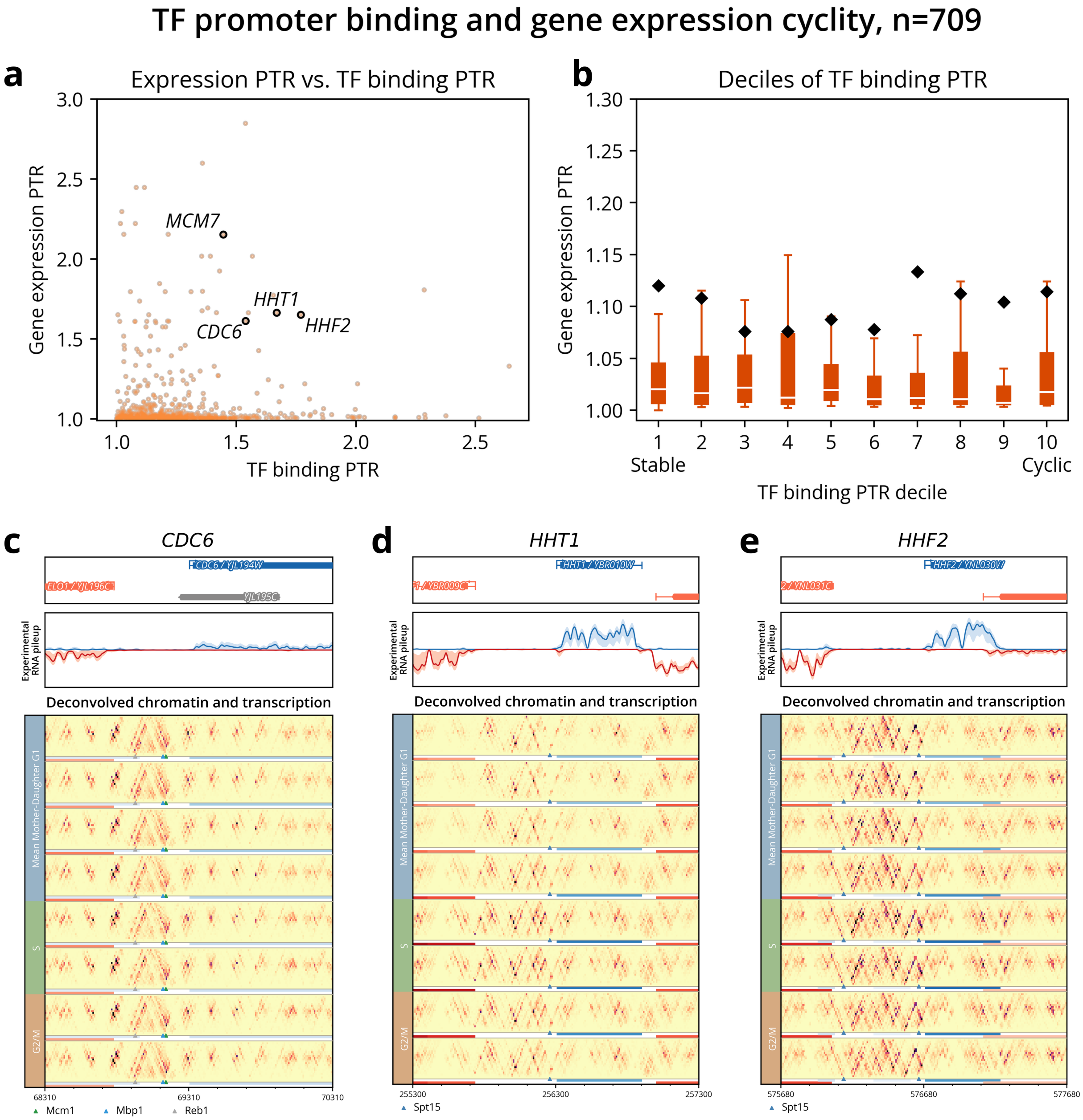


**Supplemental Figure S12**. Cyclic TF promoter binding is independent of cell cycle gene expression. **(a)** Scatter plot shows that TF binding PTR is not correlated to gene expression PTR. Genes *MCM7*, *HHT1*, *CDC6*, and *HHF2* are highlighted as notable examples, with *MCM7* previously analyzed with the *MCM2–7* complex genes. **(b)** Similar to the scatter plot findings, segmenting TF binding into deciles confirms little connection between gene expression cyclicity and TF binding cyclicity. **(c)** *CDC6* shows very subtle gene expression changes peaking in G1 (the darkest blue lines are in the top half of the deconvolved section). Meanwhile, in the promoter of *CDC6*, chromatin signal at Reb1, Mbp1, and Mcm1 sites show cyclic changes. **(d)** Histone gene *HHT1* shows dramatic cycling expression that peaks in S phase. At the binding site for general TBP factor Spt15, we note a clear concomitant signal of activation. **(e)** Likewise, at histone gene *HHF2*, a concomitant signal of activation appears at a Spt15 binding site.

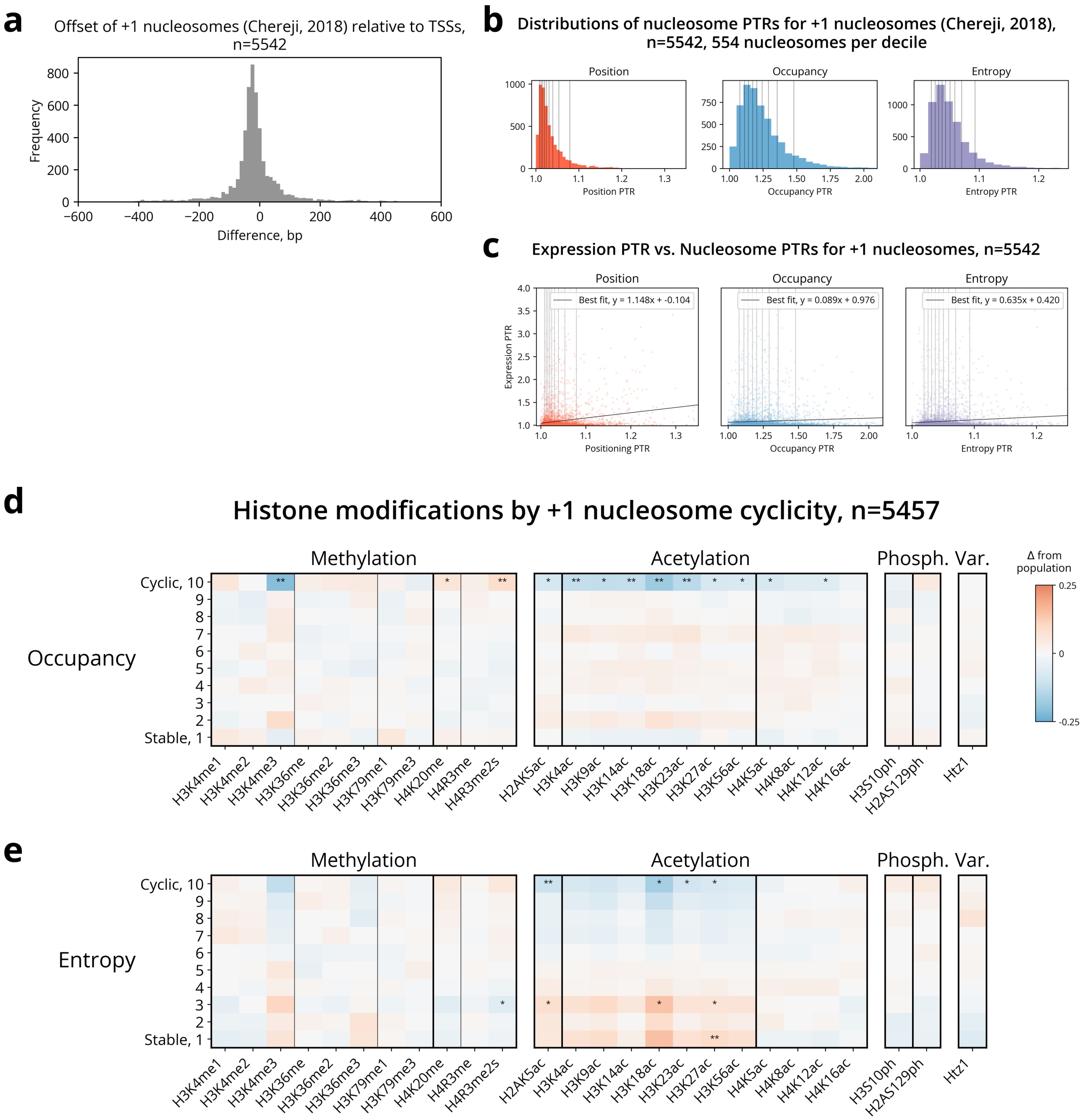


**Supplemental Figure S13**. Cyclicity of +1 nucleosomes relates to histone modification levels. **(a)** Chereji *et al.* (2018) +1 nucleosome and TSS annotations largely agree. Nearly all +1 nucleosomes are within 200 bp of annotated TSSs. **(b)** Distribution of peak-to-trough ratios (PTR) for nucleosome dynamics across three measures: (left) position, (middle) occupancy, and (right) entropy. Red lines mark the deciles for each measure. **(c)** Scatter plots show the relationship between each measure and expression. Left: +1 position cyclicity has a positive relationship with expression cyclicity, while occupancy (middle) and entropy (right) do not. **(d)** Deciles of +1 occupancy cyclicity show mostly population-level histone modifications except for the most cyclic. **(e)** Deciles of +1 entropy cyclicity mirror the observation found with position cyclicity: histone acetylation is inversely proportional to transcription cyclicity and H3K4 mono- and dimethylation reflect cyclic transcription, without a long enough period for trimethylation to occur.

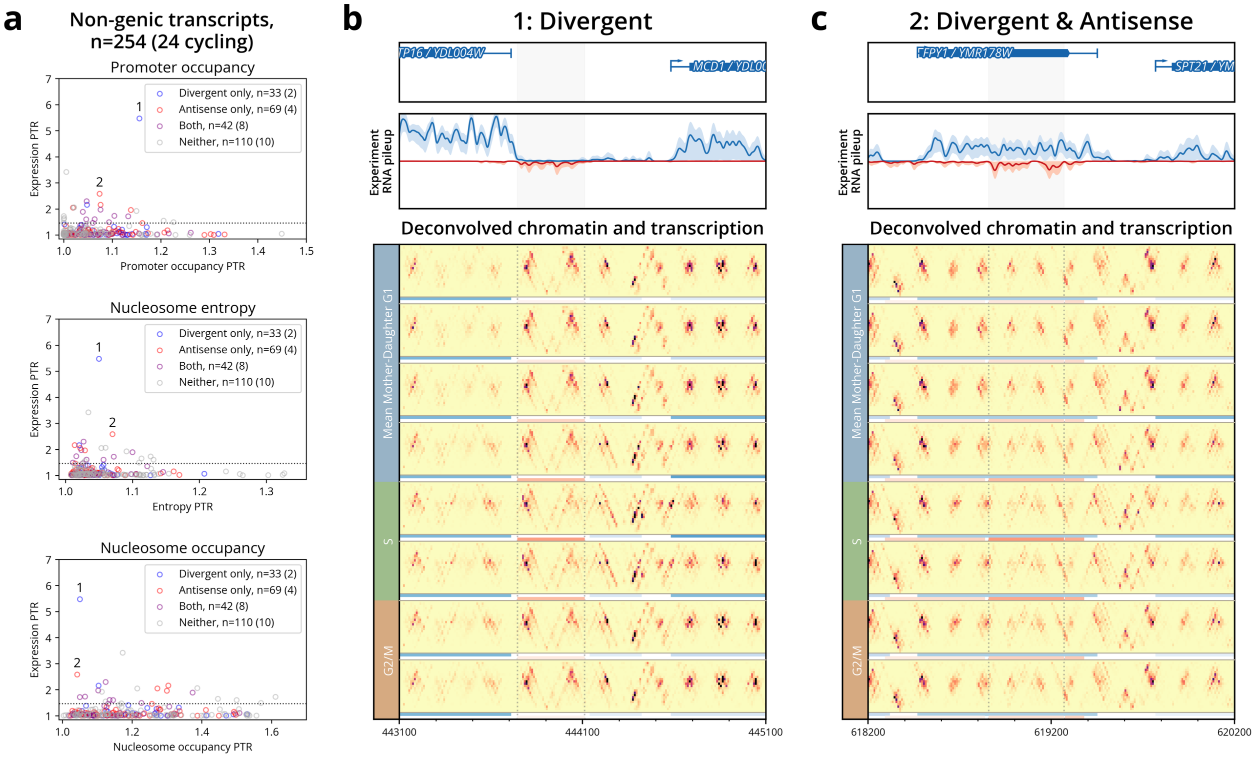


**Supplemental Figure S14**. Cell cycling non-genic transcripts activate in a variety of genomic contexts. **(a)** Scatter plots show chromatin PTR versus expression PTR for 254 non-genic transcripts, with 24 above 95th percentile threshold for cell cycle expression (colored by genomic context: divergent only, antisense only, both, or neither). While most cell cycling non-genic transcripts show subtle chromatin changes, a couple detailed examples reveal highly coordinated chromatin and transcription dynamics. **(b)** Locus-level plot shows the dynamics of a non-genic divergent transcript (region indicated by gray shading and dotted lines) sharing a promoter with S-phase gene *MCD1*. These transcripts show coordinated peak expression timing during late G1/S phase (dark red and blue lines in the middle and right ends of the deconvolved section), +1 nucleosome displacement for both transcripts, and shared promoter reorganization. **(c)** Locus-level plot shows an example of "Both" category, in which a transcript that is antisense to *FPY1* is also divergently transcribed from the promoter of *SPT21*. This transcript suggests regulation through two chromatin effects: nucleosome accessibility from stable *FPY1* transcription and shared promoter coordination with cell cycle regulated *SPT21*.

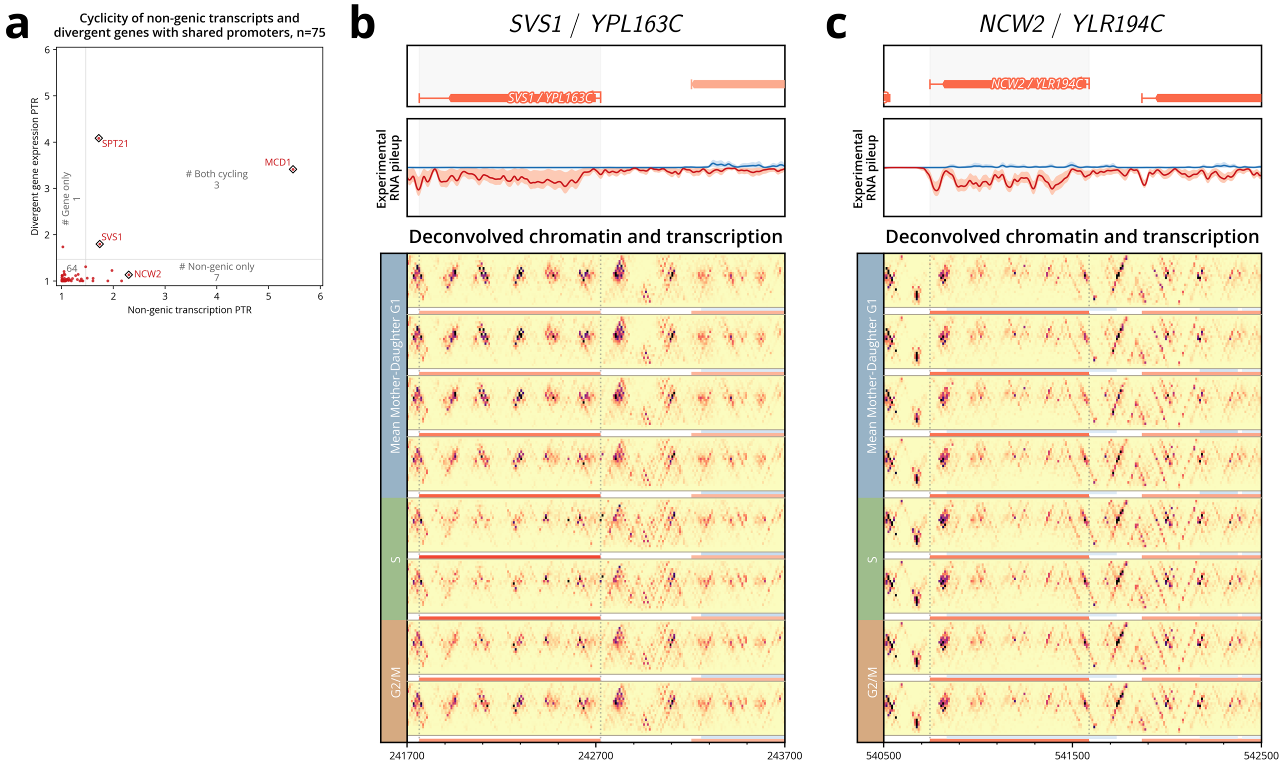


**Supplemental Figure S15**. Non-genic and divergent gene transcription rarely cycle together despite shared promoters. **(a)** Scatter plot of cycling strength (PTR) for 75 divergent gene-nongenic transcript pairs sharing promoters. Only 3 show coordinated cycling of both gene and non-genic transcript. **(b)** Cell wall gene *SVS1* represents an example where both gene and divergent non-genic transcript cycle coordinately, suggesting shared regulatory machinery. **(c)** Cell-wall remodeling gene *NCW2* exemplifies a slightly more common pattern: stable gene expression with independent cycling of the upstream non-genic transcript (light blue transcript on the far right of the plot).

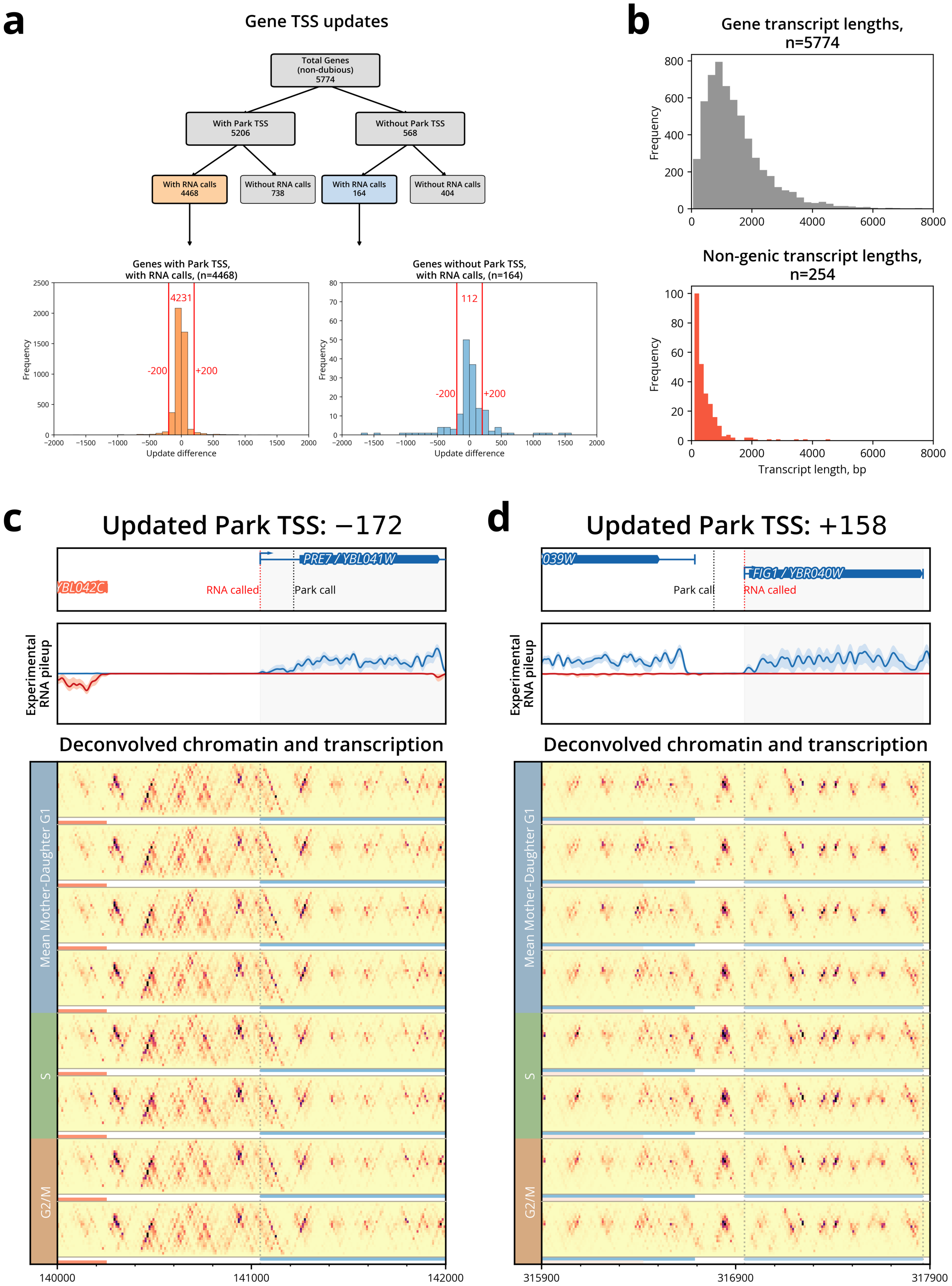
a
Gene TSS updates
Total Genes (non-dubious) 5774
With Park TSS 5206
Without Park TSS 568
With RNA calls 4468
Without RNA calls 738
With RNA calls 164
Without RNA calls 404
Genes with Park TSS, with RNA calls, (n=4468)
4231
-200
+200
Frequency
Update difference
Genes without Park TSS, with RNA calls, (n=164)
112
-200
+200
Frequency
Update difference
b
Gene transcript lengths, n=5774
Frequency
Non-genic transcript lengths, n=254
Frequency
Transcript length, bp
c
Updated Park TSS: −172
PRE7 / YBL041W
YBL042C
RNA called
Park call
Experimental RNA pileup
Deconvolved chromatin and transcription
Mean Mother-Daughter G1
S
G2/M
140000
141000
142000
d
Updated Park TSS: +158
039W
FIG1 / YBR040W
Park call
RNA called
Experimental RNA pileup
Deconvolved chromatin and transcription
Mean Mother-Daughter G1
S
G2/M
315900
316900
317900

**Supplemental Figure S16**. TSS updates using RNA-seq calls. **(a)** Flowchart showing classification of 5,774 non-dubious genes by Park TSS status and RNA-called TSS availability. 5,206 genes have Park TSS annotations and 4,468 of those have RNA-called TSSs. 568 genes lack Park TSS annotations and 164 of those have RNA-called TSSs. Histograms show the signed differences between the RNA-called TSS and the original TSS (the Park TSS for genes with such an annotation, and the 5' end of the SGD-labeled ORF for genes without such an annotation). TSSs were updated if the difference was within 200 bp, resulting in TSS updates to 4,343 genes (4,231 with Park TSSs; 112 without). **(b)** Distribution of gene (top) and non-genic (bottom) transcript lengths. Called non-genic transcripts are much shorter (most <1500 bp long) compared to genes (most <4500 bp long). **(c,d)** We show the RNA-called TSSs for two examples with sizable updates. Both the transcript boundary and the corresponding promoter architecture match the updated TSS.

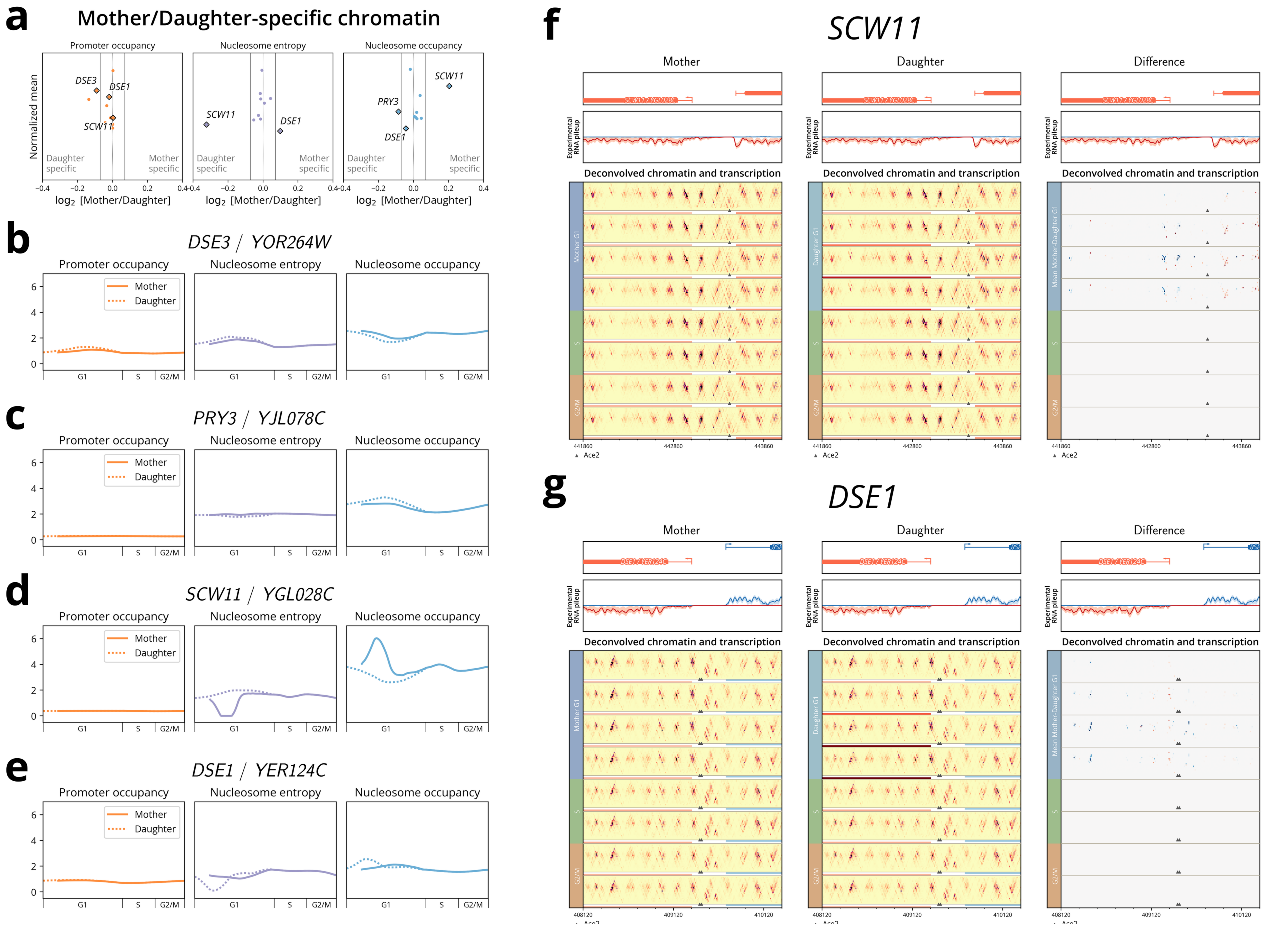


**Supplemental Figure S17**. CyCLOPS detects mother/daughter-specific chromatin differences with nucleosome-level precision. **(a)** Measures of chromatin show that most daughter-specific genes have similar chromatin signatures in both mother and daughter cells, with a few notable exceptions. Genes are plotted as $\log_2$ ratios between mother and daughter chromatin values. A threshold of $\pm \log_2 0.07$ was chosen to capture the main cluster of unchanged genes. **(b–e)** Trace plots depict differences between mother and daughter cells in G1 for promoter occupancy, nucleosome entropy, and nucleosome occupancy. **(b)** *DSE3* and **(c)** *PRY3* show virtually identical mother/daughter profiles across all three chromatin measures. In contrast, **(d)** *SCW11* and **(e)** *DSE1* display more prominent differences in nucleosome entropy and nucleosome occupancy. **(f,g)** Nucleosome-resolution heatmaps for *SCW11* and *DSE1* show mother (left), daughter (middle), and difference (right) profiles. In the difference plots, blue represents mother-specific signal and red represents daughter-specific signal. **(f)** *SCW11* shows high mother-specific nucleosome occupancy at the TSS. **(g)** *DSE1* shows enriched mother-specific occupancy at a nucleosome positioned within the gene body (a signal not fully captured by the gene body measure in **(e)**). Both examples suggest mother-specific regulatory mechanisms operating through nucleosome occupancy.

| | Replicate 1 | | Replicate 2 | |
|---|---|---|---|---|
| | **Mean** | **95%** | **Mean** | **95%** |
| $\mu_0$ | –26.790 | (–27.055, –26.474) | –20.151 | (–20.346, –19.945) |
| $\delta$ | 8.181 | (7.461, 8.819) | 10.895 | (10.437, 11.355) |
| $\sigma_0$ | 10.989 | (10.873, 11.105) | 10.631 | (10.513, 10.747) |
| $\sigma_v$ | 0.142 | (0.139, 0.145) | 0.130 | (0.127, 0.133) |
| $\lambda$ | 68.191 | (67.711, 68.691) | 60.209 | (59.870, 60.555) |
| $\gamma_1$ | 0.004 | (0.002, 0.008) | 0.002 | (0.001, 0.003) |
| $\gamma_2$ | 0.255 | (0.252, 0.258) | 0.284 | (0.282, 0.286) |
| $\mu_{\alpha 1}$ | 7.733 | (7.732, 7.734) | 7.585 | (7.584, 7.586) |
| $\sigma_{\alpha 1}$ | 0.001 | (0.001, 0.002) | 0.001 | (0.001, 0.002) |
| $\mu_{\alpha 2}$ | 1.017 | (1.001, 1.033) | 1.010 | (1.002, 1.018) |
| $\sigma_{\alpha 2}$ | 0.031 | (0.016, 0.059) | 0.016 | (0.008, 0.030) |
| halted | 0.225 | (0.220, 0.229) | 0.291 | (0.286, 0.295) |

**Supplemental Table S1**. Average and 95% confidence interval for CLOCCS fitting parameters. Parameters are defined in detail in Orlando *et al.* (2009)

### Supplemental Method S1: Sequencing data processing

#### *MNase sequencing data processing*

The MNase seq FASTQ files were aligned with the sacCer3/R64 version of the *S. cerevisiae* genome using Bowtie v1.3.1 (Langmead and Salzberg 2012) in paired end mode with the following parameters: `-n 2 -l 20 --phred33-quals -m 1 --best --strata -y`.

#### *RNA sequencing data processing*

FASTQ files of RNA-seq data were aligned to the sacCer3/R64 version of the *S. cerevisiae* genome in single-read mode using Bowtie v1.3.1 (Langmead and Salzberg 2012) with the following parameters: `-n 2 -l 20 --phred33-quals -m 1 --best --strata -y`. The RNA-seq data of the 110 min time point for Replicate 2 were discarded due to low quality.

We normalized gene transcription using transcripts per million (Wagner *et al.* 2012) for each sample in the replicate time courses.

### Supplemental Method S2: Reference data sets

#### *Gene reference data set*

Our gene set was compiled from the Saccharomyces Genome Database (SGD) using the non-dubious genes of the sacCer3/R64 genome build. This set was then filtered according to a minimum sequencing depth for both the MNase-seq for a final set of 5,578 genes.

Transcription start sites (TSS) were defined primarily according to Park *et al.* (2014). If TSS was not available, the gene boundary defined in the SGD sacCer3/R64 genome build was used.

#### *Origins of replication reference data set*

Our origins were defined as the set of 798 replication origins characterized by Belsky *et al.* (2015).

### Supplemental Method S3: Detailed replication profile optimization procedure

#### *Replication profile optimization*

Given occupancy values $\mathbf{G}_r$ and the convolution kernel $\mathbf{H}$, estimation of the replication profile matrix $\mathbf{F}_r$ proceeds as follows.

1. Initialize matrices **N** and **B**.
2. Optimize the replication profile matrix $\mathbf{F}_r$.
3. Update **N** and **B** from the current estimate of $\mathbf{F}_r$.
4. Repeat steps 2–3 until convergence.

This procedure estimates the optimal replication profile matrix $\mathbf{F}_r$ by progressively refining the normalization and baseline occupancy terms **N** and **B**. Iteration is necessary because **N** and **B** depend on $\mathbf{F}_r$, and vice versa.

#### *Initializing and updating scalar matrices **N** and **B***

Scalar matrices **N** and **B** define additional terms for normalization. **N** normalizes for the average DNA content over each sample. **B** normalizes for per-window variation.

$\mathbf{N}$ is initialized analytically from $\mathbf{H}$ and known cell cycle DNA content, using a linear approximation for S-phase that is refined during updates. The average DNA content per experimental time point is computed by summing three contributions from the columns of $\mathbf{H}$: (1) for G1 phases (recovery, mother, daughter, and halted), DNA content is 1; (2) for S phase, DNA content is linearly interpolated from 1 to 2; (3) for G2/M, DNA content is set to 2 copies. This yields $\bar{n}$, and $\mathbf{N} = \mathrm{diag}(\bar{n})^{-1}$.

$\mathbf{B}$ is initialized with the first experimental time point in $\mathbf{G}_r$ for each replicate. These time points provide an initial estimation of the single DNA copy occupancy for each genomic window.

Once $\mathbf{F}_r$ is available, both matrices are updated using the replication profile optimization equation:

$$\mathbf{G}_r = \mathbf{N}\mathbf{H}\mathbf{F}_r\mathbf{B}$$

$\mathbf{N}^*$ is updated using $\mathbf{F}_r$ directly, replacing the linear S-phase approximation with the estimated replication profile. The element-wise ratio of $\mathbf{G}_r\mathbf{B}^{-1}$ and $\mathbf{H}\mathbf{F}_r$ is computed, and its row means yield the updated diagonal of $\mathbf{N}^*$. Also note that $\bar{n}^*$ is normalized by the number of deconvolved time points $u$:

$$\bar{n}^* = \left(\mathbf{G}_r\mathbf{B}^{-1} \oslash \mathbf{H}\mathbf{F}_r\right) \frac{1}{u}\mathbf{1}_{(u\times 1)} \qquad \text{(Row means of element-wise ratio)}$$
$$\mathbf{N}^* = \mathrm{diag}(\bar{n}^*)^{-1}$$

Where $\oslash$ denotes element-wise division.

$\mathbf{B}^*$ is set to the ratio of $\mathbf{G}_r$ to the prediction term prior to $\mathbf{B}$ scaling, $\mathbf{N}\mathbf{H}\mathbf{F}_r$. The column sums of each are computed, and their element-wise division yields the updated diagonal:

$$b_{\text{diag}} = \left(\mathbf{G}_r{}^{\top}\mathbf{1}_{(m\times 1)}\right) \oslash \left[(\mathbf{N}\mathbf{H}\mathbf{F}_r)^{\top}\mathbf{1}_{(m\times 1)}\right] \qquad \text{(Element-wise division of column sums)}$$
$$\mathbf{B}^* = \mathrm{diag}(b_{\text{diag}}) \qquad \text{(Construct a diagonal matrix)}$$

### Supplemental Method S4: Chromatin deconvolution model details

#### *Normalization of MNase-seq for chromatin deconvolution*

For proper analysis of cell cycle chromatin, MNase-seq data must be normalized to correct for MNase digestion variability while preserving copy number variation for downstream correction. Digestion variability between samples creates differences in fragment length distributions that can confound downstream analysis by artificially inflating/deflating nucleosome or subnucleosome occupancy change. At the same time, normalization must preserve the natural copy number variation arising from replication timing, as this variation is essential for downstream copy number correction.

Prior to deconvolution, the genome is segmented into 10 kb windows for computational efficiency. For each window represented as 2D histograms (Fig. 1B), fragment length normalization is applied to correct for MNase digestion variability while preserving copy number variation. Each window is scaled to match a target distribution computed globally across all samples.

Additionally, a per-window scaling factor is applied that preserves the natural copy number variation over time of that genomic region. This two-part scaling approach ensures consistent fragment length distributions across samples while maintaining the overall occupancy variation between windows needed for downstream copy number correction.

#### *Construction and smoothing of idealized average single-cell profiles $\mathbf{F}_c$*

Following Guo *et al.* (2013), the deconvolved average single-cell profiles $\mathbf{F}_c$ were composed of discrete blocks of time points that corresponded to cell cycle phases within the branching process model:

$$\mathbf{F}_c = [\mathbf{F}_{RG1}, \mathbf{F}_{MG1}, \mathbf{F}_{DG1}, \mathbf{F}_S, \mathbf{F}_{G2M}]$$

The blocks comprising the initial recovery, mother, and daughter branches of the model are then concatenated in order to apply smoothing regularization (as described in the manuscript):

$$\mathbf{F}_{\text{Recovery}} = [\mathbf{F}_{\text{RG1}}\mathbf{F}_{S}\mathbf{F}_{G2M}]$$
$$\mathbf{F}_{\text{Mother}} = [\mathbf{F}_{MG1}\mathbf{F}_{S}\mathbf{F}_{G2M}]$$
$$\mathbf{F}_{\text{Daughter}} = [\mathbf{F}_{DG1}\mathbf{F}_{S}\mathbf{F}_{G2M}]$$

The total number of indices (i.e., deconvolved time points) in each of these branches was chosen to be a power of two (we used 128) to allow for simplicity in generating the Symlet wavelets. Each of $\mathbf{F}_{RG1}$, $\mathbf{F}_{MG1}$, and $\mathbf{F}_{DG1}$ was allocated 62 indices, while 66 indices were given to the post-G1 phases ($\mathbf{F}_{S}$ and $\mathbf{F}_{G2M}$), ensuring a sufficient number of indices for sub-minute temporal resolution after deconvolution. Edge effects were handled by duplicating the mother and daughter branches and by mirroring the start of the recovery branch, respectively.

### Supplemental Method S5: Selection of chromatin deconvolution regularization parameters

*Selection of smoothness regularization parameter $\gamma$*

The regularization parameter $\gamma$ controls the degree of smoothness imposed on the deconvolved average single-cell chromatin profiles, ensuring biologically plausible cell cycle transitions.

To determine the optimal $\gamma$ value, we deconvolved one hundred randomly selected genomic windows of 1000 bp. For each window, we identified the knee point where increasing $\gamma$ produced diminishing returns in improving solution smoothness (Satopaa *et al.* 2011). The optimal $\gamma$ value was determined as the average of these knee points across all windows.

*Selection of MG1/DG1 differences regularization parameter $\kappa$*

Due to $\alpha$-factor arrest which occurs late in G1, the first cell cycle's G1 is treated separately as a recovery G1 (RG1), distinct from later cell cycles which contain equal numbers of cells undergoing mother G1 (MG1) and daughter G1 (DG1). Without information provided by the initial RG1, these latter cell cycles may present model identifiability issues for distinguishing between MG1 and DG1 reads.

The model was therefore regularized using an L1 norm to limit the amount of difference between MG1 and DG1. To determine the proper weight for this regularization term, $\kappa$, a conservative threshold of 5% was set as the amount of chromatin in the genome that is considered MG1- or DG1-specific. One hundred random genome regions of 1000 bp were sampled to identify the $\kappa$ value where this threshold is met. The average optimal $\kappa$ for these one hundred regions was chosen as the optimal $\kappa$ value for genome-wide chromatin deconvolution.

*Selection of halted cells regularization parameter $\eta$*

In the cell cycle experiment, halted cells are expected to be similar to recovery G1 (RG1) cells. A regularization term $\eta$ was applied to constrain the difference between the halted cell chromatin occupancy profiles and the average RG1 profile, ensuring halted cells remain sufficiently similar to RG1 profiles.

To determine the optimal $\eta$ value, we deconvolved one hundred randomly selected genomic windows of 1000 bp each. For each window, we identified the knee point where increasing $\eta$ produced diminishing returns in reducing RG1-halted differences (Satopaa *et al.* 2011). The optimal $\eta$ value was then determined as the average of these knee points across all windows.

### Supplemental Method S6: Transcription deconvolution model

Deconvolution of gene expression was performed on the set of 5,578 genes and 171 non-genic transcripts using the deconvolution model described for the chromatin with a single vector of values to deconvolve ($v = 1$).

Selection of regularization parameter $\gamma$ was computed for every transcript individually, as previously described in Guo *et al.* (2013).

### Supplemental Method S7: Selection of parameters estimating the delay between cytokinesis and cell wall breakdown, $\alpha_1, \alpha_2$

Flow cytometry data are limited in capturing the timing of mitosis, as they may misclassify two post-cytokinesis G1 cells as a single G2 cell when the cell wall remains intact between two newly divided cells. Estimating the delay parameters $\alpha_1$ and $\alpha_2$ (a separate one for each replicate) between cytokinesis and cell wall breakdown is therefore critical for the deconvolution model to properly distinguish the end of M phase from the start of G1.

To determine optimal $\alpha$ values for each replicate, we used the expression of daughter-specific genes previously characterized by Colman-Lerner *et al.* (2001) and utilized by Guo *et al.* (2013): *DSE1*, *DSE2*, *DSE3*, *DSE4*, *ASH1*, *EGT2*, *AMN1*, *PRY3*, *SCW11*, and *CTS1*. These genes are specifically upregulated in DG1 following mitosis and are therefore indicative of proper $\alpha$ parameter selection.

The selection of $\alpha$ values was constrained by two criteria. First, $\alpha$ values must be sufficiently long for daughter-specific genes to be primarily expressed in DG1. Second, the selected $\alpha$ values must result in highly correlated daughter-specific expression profiles between the two replicates after deconvolution. Because these individually-determined parameters are subsequently used together in the combined deconvolution model, this constraint ensures the combined model produces stable results. Values of $\alpha_1 = 34$ minutes and $\alpha_2 = 28$ minutes were selected as they satisfied both constraints.

### Supplemental Method S8: Software and tools

For all deconvolution runs, the convex optimization library CVXPY was used (Agrawal *et al.* 2018; Diamond and Boyd 2016). For efficiency the solver MOSEK (MOSEK ApS 2025) was used for all long running deconvolution runs. The default CVXPY solver Clarabel (Goulart and Chen 2024) was also run to confirm consistency between solvers.
